# Treatment Resistant Persister Cells Exploit Macrophage Lipid Metabolism to Sustain Glioblastoma Growth

**DOI:** 10.1101/2025.06.07.658345

**Authors:** Avirup Chakraborty, Changlin Yang, Aryeh Silver, Diana Feier, Guimei Tian, Avinash Pittu, Christina Von Roemeling, Nagheme Thomas, Michael Andrews, Cole Conforti, Olusegun O. Sobanjo, Nyla T. Searl, Miruna N. Anica, Raquel McTiernan, Maryam Rahman, Jianping Huang, Jeffrey K. Harrison, Duane A. Mitchell, Matthew Sarkisian, Loic P. Deleyrolle

## Abstract

Glioblastoma (GBM) displays pronounced intratumoral heterogeneity, posing significant challenges to understanding its biology and developing effective treatments. Using spatial multi-omics, *in vivo* functional assays, and systems-level analysis, we delineate the diverse metabolic and immune architecture of GBM. We identify a lipid-dependent lineage of treatment-resistant persister cells (TRPCs) that engage tumor-associated macrophages (TAMs) in a spatially organized, metabolically specialized crosstalk. TRPCs co-opt CCR2⁺, CSF1R⁺, CD163^+^ TAMs for lipid scavenging and acquisition, promoting a pro-tumorigenic and immunosuppressive microenvironment. This cooperative axis is critically dependent on lipid chaperones like FABPs, whose targeting disrupts TAM recruitment, remodels immune composition, and suppresses tumor growth. Retrospective clinical analyses reveal that elevated TRPC-associated transcriptome may serve as stratification criteria to identify patients benefiting from lipid-lowering therapies like statins. Our findings uncover a targetable immunometabolic circuit between TRPCs and TAMs and support the development of precision therapies that disrupt lipid-fueled tumor-immune cooperation in GBM.

**In brief:** Treatment-resistant persister cells (TRPCs) in glioblastoma spatially engage TAMs to facilitate lipid transfer, thereby sustaining tumor growth and promoting immune evasion. Targeting this TRPC–TAM metabolic axis reprograms the immunosuppressive microenvironment and improves therapeutic outcomes, revealing a clinically actionable metabolic vulnerability with potential for precision immune-metabolic interventions in GBM.

**Highlights:**

- GBM exhibits spatially resolved heterogeneity revealing correlative metabolic and immune micro-niches
- TRPC lineage promotes a pro-tumorigenic and immunosuppressive microenvironment by recruiting lipid-specialized TAMs
- TRPC lineage hijacks TAMs for metabolic support via stimulating lipid transfer and acquisition
- Disruption of the TRPC-TAM lipid axis, including through FABP3 targeting, reprograms the immune landscape, limits TAM recruitment, and impairs tumor progression
- TRPC lineage and associated micro-niche transcriptomic profiles can serve as criteria for patient stratification and identification of responders to lipid-lowering therapies, such as statins

## INTRODUCTION

Glioblastoma (GBM) is the most aggressive and common form of primary brain tumor in adult and is highly refractory to therapy due to its robust inter- and intra-tumoral heterogeneity. This complexity spans genetic, cellular, metabolic, and immune variations, which manifests in distinct spatial niches within the tumor microenvironment (TME) ^1–9^. This diversity and plasticity are not only central to therapeutic resistance and tumor recurrence but also pose a significant challenge to implementing universal treatment strategies. In our efforts to decipher tumor heterogeneity, our group identified a drug-tolerant, aggressive cell phenotype originating from a subpopulation of slow-cycling cells (a.k.a SCCs) ^10–12^. These cells give rise to and maintain a lineage of cells we herein refer to as treatment-resistant persister cells (TRPCs), which we previously showed preferentially rely on lipid metabolism ^11^ ^12^. In contrast, treatment-sensitive cells (TSCs), also previously mentioned as Fast Cycling Cells (FCCs), exhibit an upregulated aerobic glycolysis pathway ^10^ ^12,13^. In this study, we employed an orthogonal multi-omics strategy combined with *in vivo* functional assays and advanced network analysis to reveal the in-depth molecular landscape of the metabolically heterogenic cellular micro-niches in a treatment-naïve murine model of GBM and human tissue samples. We investigated the spatial interactions between GBM cells and the immune compartment to assess how metabolic heterogeneity influences immune diversity, specification, and intercellular communication. This led to the identification of enriched molecular and biological pathways that drive spatially distinct recruitment of TAMs, which metabolically support TRPCs via lipid transfer, enabling them to overcome the metabolic pressures of the TME and sustain their aggressive, recurrence-driving phenotype. We assessed this TRPC-TAM homeostasis as targetable vulnerability and validated its clinical relevance through a retrospective study. Patients with a high TRPC transcriptional signature demonstrated improved responses to lipid-lowering therapies such as statins, which not only enhanced survival but also induced transcriptional remodeling linked to increased immune responsiveness, highlighting a potential synergy with immune checkpoint inhibitors.

In summary, our findings reveal the metabolic interplay between TRPCs and TAMs as a fundamental mechanism driving GBM biology. Targeting the lipid-driven TRPC-TAM axis offers a promising therapeutic avenue, especially in combination with immunotherapies, and may pave the way for personalized treatment strategies based on metabolic and immune profiling.

## RESULTS

### Tumor heterogeneity encompasses immune heterogeneity in murine GBM

Our previous research highlighted metabolic diversity within the TME, showing that a lineage of TRPCs in human GBM tumors preferentially utilize lipid metabolism rather than aerobic glycolysis mostly observed in TSCs ^10,11,13,14^. TRPCs exhibit aggressive tumor-initiating properties, increased invasion, and resistance to therapy ^10,11,13,14^. To investigate the interactions between TRPCs and immune cells, and evaluate the correlation between metabolic and immune heterogeneity, we utilized an immunocompetent system with the KR158 murine glioma model ^15^. We demonstrated this model replicates the key characteristics of this lineage observed in human GBM ^12^. Our previous studies comparing the growth of tumors generated by the intracranial transplant of total unsorted KR158, TSCs, or TRPCs in immune competent mice revealed that TRPC-derived tumors showed a faster progression compared to the other groups, resulting in shorter survival time ^12^. However, using immunocompromised NSG mice, TRPCs did not retain their growth advantage over TSCs. Instead, both groups exhibited similar tumor progression and survival rates (***Fig. 1A, B***). These findings suggest that the immune compartment plays distinct regulatory roles in modulating the growth of TRPC and TSC lineages, thereby influencing overall tumor progression. Based on these differences, we hypothesized that tumor cell heterogeneity could shape the diversity of the immune landscape within the GBM microenvironment and that immune infiltrates may play a specific supportive role, specifically in TRPC-driven tumor progression. To test this, the transcriptomic profile of CD45^+^ cells sorted from TSC- and TRPC-derived tumors were interrogated via RNA sequencing (***Fig. S1A***). Principal component analysis identified specific projection patterns indicating divergent immune landscape between the groups (***Fig. 1C*, *Table S1***). The expression level of *67 immune gene signatures* identifying 9 different immune cell populations, was represented as a heatmap and revealed a dichotomous immune gene regulation between the different tumors (***Fig. S1B***). Pathway enrichment analysis revealed an overall upregulation of myeloid-associated genes and a concurrent downregulation of lymphoid-associated genes in TRPC tumors (***Fig. 1D***). In contrast, TSC tumors exhibited the opposite trend (***Fig. 1D-F*, *Table S2***). Flow cytometry-based phenotypic analysis of CD45^+^ immune infiltrates further confirmed the positive regulation of T cells in TSC tumors, showing significantly higher T cell infiltration (***Fig. 1G*, *S1C***). More specifically, the immune infiltrates observed in tumors originating from TSCs exhibited a distinctive gene expression pattern associated with the upregulation of adaptive anti-tumor immune response pathways (***Fig. 1H***). This pattern was further substantiated by elevated TCR signaling, cytokine signaling, and expression of genes regulating T cell memory and helper T cell differentiation within the immune infiltrates of TSC tumors (***Fig. 1H***). We performed TCR profiling to analyze clonotype amplification, and unique rearrangements of the J and V segments of TCR β chain from CD45^+^ tumor-infiltrating cells which are indicators of immune activation. Results showed a higher frequency of dominant clonotypes in TSC tumors (***Fig. 1I-J*, *S1D***), with the most expanded clonotype (TRBV5/TRBJ2-1) comprising 30.8% of all clones (***Fig. 1K***). In contrast, TRPC infiltrates had a more heterogeneous TCR profile, with the dominant clonotype only representing 8.3% of all clones. T-cell activation and immune responses were evaluated *in vivo* by measuring IFNγ expression and secretion within the TME. KR158 cells expressing luciferase (KLuc) were delivered intracranially in the interferon-Gamma Reporter with Endogenous polyA Transcript (GREAT) mice. Bioluminescence combined with fluorescence *in vivo* imaging revealed that greater volumes of TRPC-derived tumors were associated with lower levels of IFNγ (***Fig. 1L, M, and* *S1E***), confirmed by ELISA and RNA sequencing (***Fig. 1N, O***). Together, these studies establish that tumors derived from TSCs present higher immunogenicity with a more inflamed environment and enhanced T cell response compared to TRPCs. To functionally evaluate the immunogenicity of TSCs *in vivo* and to demonstrate that lymphocytes mediate the control of TSC tumors in an intact immune system we implanted TSCs and TRPCs into the brains of Fox Chase SCID mice, which are deficient in both T and B cells, and CD8-Knock Out (KO) mice, which specifically lack CD8+ T cells. In both contexts, TSCs exhibited more aggressive and rapidly progressing tumors, leading to shorter survival times (***Fig. 1P-R***). This suggests that with an intact immune system, CD8 T cells can target TSCs and suppress tumor growth, whereas TRPCs evade immunosurveillance, resulting in more aggressive tumors ^12^. The diminished adaptive anti-tumor immunity observed in TRPC tumors, may be attributed to greater recruitment of immunosuppressive tumor-associated myeloid cells (TAMCs), including macrophages and myeloid-derived suppressor cells (MDSCs). This hypothesis is supported by the upregulation of genes associated with innate immunity, myeloid-mediated immune evasion, and TLR signaling (***Fig. 1S*).** Enrichment of TAM signature, upregulation of gene sets indicative of bone marrow-derived macrophages (BMDMs), and polarization towards M2-like macrophages were also observed in TRPC tumor immune infiltrates (***Fig. 1T-V*, *S1F-L, Table S3***). Overall, our data confirms a divergent immune composition and status within the TME shaped by TRPCs and TSCs and suggests that a key difference between these two populations of GBM cells may lie in their distinct interactions with the immune system.

**Figure 1:**
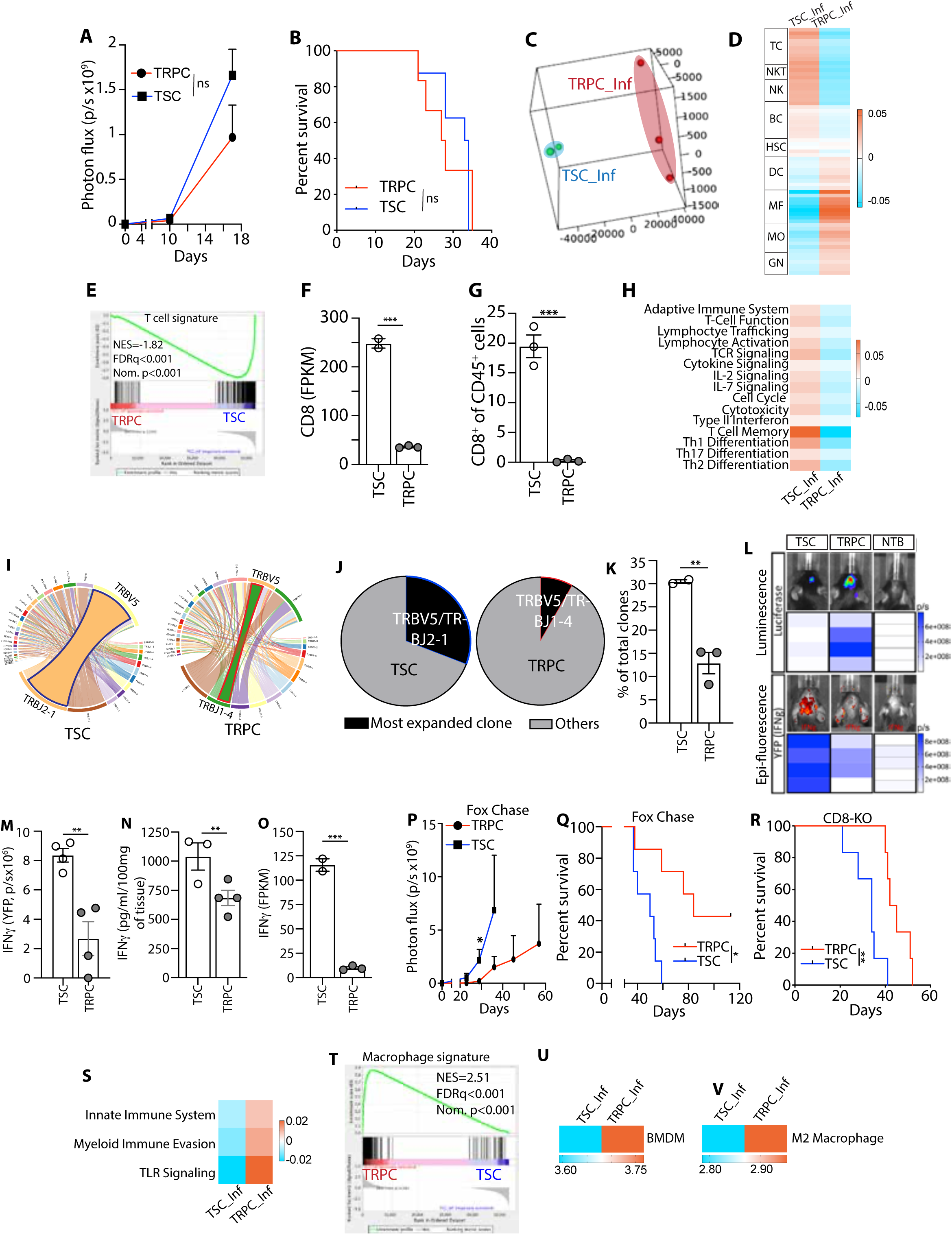
Tumor cell lineages associated with specific immune contexture. **A)** Bioluminescence-based monitoring of murine (m) TRPC/mTSC tumor growth in NSG mice (n=6-8; two-way ANOVA) **B)** Kaplan-Meier survival in NSG mice. (n=6, 8; Log-rank test) **C)** 3D PCA analysis of immune infiltrates (ImmInfs) from mTRPC and mTSC derived tumors grown in immunocompetent C57BL/6 mice **D)** Mean Heatmap presents distinct level of expression of 9 gene signatures [T cells (TC), natural killer T cells (NKT), Natural killer cells (NK), B cells (BC), hematopoietic stem cells (HSC), dendritic cells (DC), macrophages (MF), monocytes (MO), granulocytes (GN)] from RNA sequencing data. **E)** GSEA of T cell gene signature in mTRPC/mTSC tumors **F)** FPKM values assessing CD8 expression in mTRPC/mTSC tumor ImmInfs (Unpaired t-test) **G)** Flow cytometric analysis CD8+T cells within CD45+ cell population of mTRPC/mTSC tumor ImmInfs. (Unpaired t-test) **H)** Heatmaps showing adaptive immune response pathways of ImmInfs in mTRPC/mTSC tumors **I)** Circos plot comparing amplification of TCRβ chain V/J segment in mTRPC/mTSC tumor ImmInfs **J)** Pie chart showing the dominant TCRβ clonotype frequency **K)** Bar diagram of J (Unpaired t-test) **L-O)** *In vivo* expression of IFN-γ in mTRPC/mTSC tumor-bearing and no tumor control (NTC) GREAT mice **(L)** Tumor growth and IFN-γ expression **(M)** Bar diagram: cranial YFP intensity (IFN-γ reporter) by epifluorescence (Unpaired t-test) **(N)** IFN-γ levels in tumor homogenates (Unpaired t-test) **(O)** FPKM values for IFN-γ expression (Unpaired t-test) **P-Q)** (P) Bioluminescence-based monitoring of mTRPC/mTSC tumor growth (n=7,7; two-way ANOVA); and (Q) Kaplan-Meier survival in Fox Chase SCID mice (n=7,7; Log-rank test) **R)** Kaplan-Meier survival of mTRPC/mTSC tumor-bearing CD8KO (n=6,6; Log-rank test) **S)** Mean heatmaps of innate immune response pathways of ImmInfs in mTRPC/mTSC-derived brain tumors. **T)** GSEA of Macrophage gene signature in mTRPC/mTSC tumor ImmInfs **U)** Mean Heatmap of BMDM in mTRPC/mTSC tumors ImmInfs **V)** Mean Heatmap of M2 macrophage in mTRPC/mTSC tumors ImmInfs

### Geospatial immune profiling uncovers intratumoral heterogeneity with differential immune recruitment within TRPC and TSC niches

To dissect the unique characteristics and niche-shaping capabilities of distinct glioma cell populations, the TRPC and TSC lineages were initially studied separately. While this approach allowed us to delineate lineage-specific features, it does not fully capture the complexity of the human disease, where both lineages coexist within the same tumor. To address this limitation and better model the intratumoral heterogeneity, we performed high-resolution spatial profiling of immune cell infiltration within tumors harboring both TRPC and TSC populations using the GeoMx Digital Spatial Profiler (DSP). This geospatial analysis enabled a more physiologically relevant understanding of lineage-specific immune interactions within the native tumor context. GFP-tagged TRPCs and RFP-tagged TSCs were co-transplanted intracranially. GFP, RFP and CD45 labeling were used as morphology markers to define regions of interest (ROIs) enriched in TRPCs or TSCs from which probes from CD45 positive cells were collected (***Fig. 2A***). Pre-tagging of the tumor cell populations allowed us to perform lineage tracing and geospatial distribution of their respective progenies. 3D PCA demonstrated a differential clustering of CD45^+^ cells (***Fig. 2B***). Similar to the results observed in ***Figure 1***, these experiments revealed distinct immune micro-niches within the TRPC and TSC regions of the same tumor. The heatmap depicts the level of immune probes detected in the distinct lineage-specific ROIs (***Fig. 2C***) highlighting the dichotomous immune contextures between the two TMEs, where TRPC-enriched regions show a greater presence of TAMCs but decreased levels of CD8^+^ T lymphocytes compared to TSC-enriched regions (***Fig. 2D-F***). Notably, we also identified a reduced presence in TRPC niches of CD34^+^ hematopoietic stem cells (HSCs), which have been shown to enhance tumor-specific T cell activation (***Fig. 2G***) ^16^.

**Figure 2:**
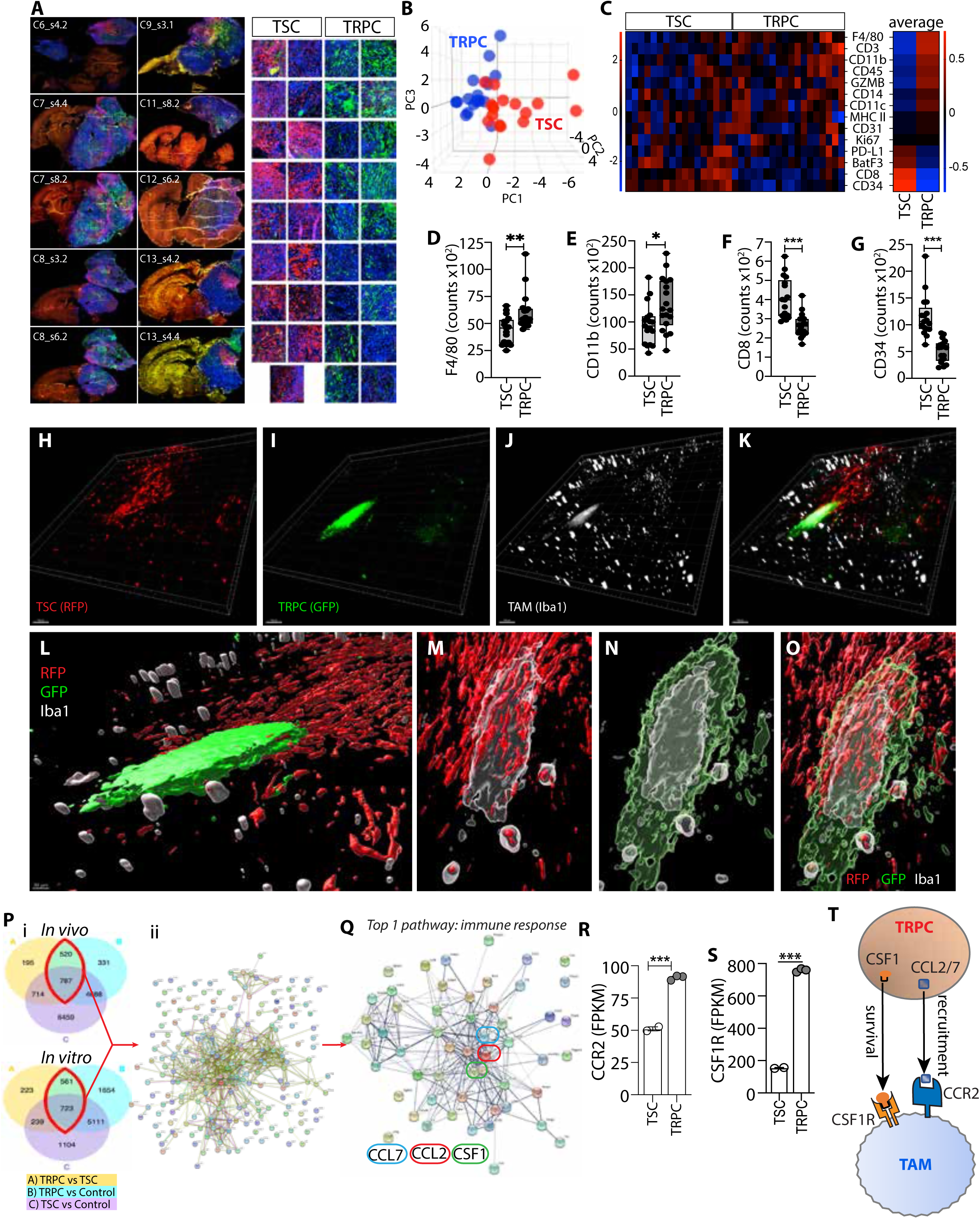
Distinct immune recruitment in mTRPC and mTSC Niches. **A)** Immunofluorescence staining of murine brain sections (n=10, see methods for section coding system) using morphological markers (RFP, GFP, CD45) to identify ROIs (n=35) **B)** 3D PCA proteomic analysis of mTRPC/mTSC tumor ImmInfs (n=17, 18) **C)** Heatmap of immune probe expression within mTRPC/mTSC ROIs (n=17, 18) **D-G)** Bar diagram (C) of ImmInfs (D) F4/80+ (E) CD11d+ (F) CD8+ and (G) CD34+ cells; (n=17, 18, Unpaired t-test) **H-K)** 3D immunofluorescence images of mTME showing (H) TSC region, (I) TRPC regions, (J) TAMs, (K) Composite image (representative of 4 different analyzed brains) **L-O)** 3D rendered images of the mTME. (L) TME showing RFP+TSC and GFP+TRPC region, (M) TAMs and TSCs, (N) TAMs and TRPCs, (O) Combined channels (representative of 4 different brains analyzed) **P (i)** Van-diagrams of differentially regulated genes between mTRPCs, mTSCs, and non-tumor control; **(ii)** Differentially up-regulated genes (n=3) **Q)** Top differentially upregulated pathway in mTRPC (n=3) **R-S)** FPKM values for CCR2 (R) and CSF1R (S) in ImmInfs; (Unpaired t-test) **T)** Schematic diagram showing signaling supporting TAM recruitment (CCL2/7-CCR2) and survival (CSF1-CSF1R).

To further our understanding of TAM distribution within the TME, we utilized 3D reconstruction for detailed spatial visualization. Our analysis confirms a preferential accumulation of TAMs in the TRPC regions even within heterogeneous micro-niches where TSC and TRPC regions were in proximity (***Fig. 2H-O****, video S1*), potentially supporting a spatially organized interaction between immune cells and TRPCs. This further supports the notion that TRPCs play a pivotal role in recruiting and retaining TAMs, acting as focal points for their accumulation. This process ultimately contributes to the formation of a dichotomous TAM-rich immune contexture, as illustrated in *Figure 1*. This differential distribution may result from the distinct upregulation of specific molecular signals or enriched pathways in the TRPC regions that promote TAM recruitment, survival, or activation. To identify the potential molecular pathways, TRPCs and TSCs were isolated via flow cytometry from both *in vitro* cultures and tumors generated by each cell population, followed by RNA sequencing to compare their transcriptional profiles (***Fig. 2P***). Gene expression analysis identified a distinct gene network comprising 228 genes that were commonly enriched in TRPCs and in TRPC-derived tumors, (***Fig. 2P***, *Table S4*). Gene Ontology analysis revealed that the most differentially regulated pathway associated with this network is linked to modulating immune processes (***Fig. 2Q***; *Table S5*). These findings suggest that TRPCs and TSCs activate distinct transcriptional programs that regulate tumor-immune interactions and immune activation. Interestingly, CCL2, CCL7, and CSF1 were identified as central nodes within this gene network, suggesting their potential role as key regulators of the immune response pathway (***Fig. 2Q***). Both CCL2 and CCL7 serve as ligands for the chemokine receptor CCR2 ^17^, which was found to be overexpressed in TRPC tumor infiltrates (***Fig. 2R***). Previous studies, including our own, have established the CCR2 pathway as a critical mediator of immunosuppressive myeloid cell trafficking into the glioma microenvironment ^16,18,19^ ^20–26^. Notably, circulating levels of CCL2 and CCL7 in the serum were significantly elevated in mice bearing TRPC tumors compared to those implanted with TSCs, indicating also systemic immunological differences between the groups (*Fig. S2A, B*). Additionally, colony-stimulating factor 1 (CSF1), plays a well-documented role in supporting the survival and proliferation of TAMs, thereby promoting tumor progression and suppressing anti-tumor immune responses ^27–30^. TRPC tumor immune infiltrates exhibited increased expression of CSF1R compared to the TSC group (***Fig. 2S***). Based on these findings, we propose, and examine further below that TRPCs actively contribute to establishing an immunosuppressive TME by driving the recruitment and maintenance of TAMs through the CCR2 and CSF1R canonical pathways (***Fig. 2T***).

### Lipid-rich areas associate with TRPCs and immune infiltrates over expressing lipid trafficking regulators

Our previous research indicated that GBM TRPCs exhibit unique metabolic properties, relying strongly on lipid metabolism for energy storage and utilization ^11^. The current study found that the TME created by these lipophilic TRPCs is enriched in immunosuppressive TAMs (***Fig. 1*, *2***), which can exhibit specific metabolic characteristics specializing them in lipid transport and exchange ^31,32^. Supporting these studies, our data showed that TRPC TAMs overexpressed proteins involved in lipid trafficking and metabolism (***Fig. 3***). Gene expression analysis revealed ApoE, a lipid-transporting **protein** ^33–36^ to be overexpressed in TRPC tumor immune infiltrates by several orders of magnitude when compared to other genes and compared to TSC infiltrates and normal brain ^37^ (***Fig. 3A***). TRPC immune infiltrates also up-regulated lipid effluxing and apolipoprotein lipidation genes ABCA1 and ABCG1 (***Fig. 3B, C***) ^38–43^. Similar observations were made in human GBM patients, where these regulators of lipid metabolism were overexpressed in GBM tissue (***Fig. 3D-F***). Furthermore, these genes were found to be positively correlated with each other suggesting a coordinated regulation of lipid metabolism pathways (***Fig. 3G*, *S3A-B***). After establishing differential expression of lipid metabolism genes, we investigated the cellular distribution of the corresponding proteins in the murine TRPC tumor tissue network. Immunohistochemistry revealed the presence of TAMs expressing APOE and ABCG1 within the TRPC TME (***Fig. 3H-J***). APOE, ABCA1, and ABCG1 proteins were also found in human GBM specimens (***Fig. S3C-E***), and like in our murine model, significant positive correlations between these lipid trafficking genes and different markers of macrophages and MDSCs were identified in GBM patients (***Fig 3K*, *L, S3F-I***). Our previous study demonstrated that human GBM TRPC exhibit enhanced lipid metabolism mediated by FABP7 and FABP3 ^11^. These proteins are essential for the transport and storage of fatty acids ^11,44–52^. Similarly, we found that TRPCs in our murine model overexpress FABP3 (***Fig. 3M***). Immunohistochemistry revealed a strong expression of FABP3 in tumor cells (***Fig. 3N***), located in proximity to ABCG1^+^ cells (***Fig. 3O***). Additionally, Oil Red O staining ^53–55^ demonstrated the significant presence of lipids in TRPC tumors (***Fig. 3P***). To further understand the spatial distribution of lipids within the TME, we stained the lipid droplets in murine brain tumors containing both GFP-labeled TRPCs and RFP-labeled TSCs. 3D reconstruction demonstrated preferential accumulation of lipid droplets in TRPC-enriched regions compared to TSC regions (***Fig. 3Q*, *Video S2)***. This indicates that these regions may assist TRPCs in fulfilling their metabolic requirements, reflecting their metabolic reprogramming. High magnification imaging revealed a significant amount of lipid deposits in direct contact with and surrounding TRPCs (***Fig. 3R*, *S3J***). The application of a transparency filter confirmed the presence of intracellular lipids within the TRPCs (***Fig. 3S***). Finally, ***Figure 3T*** illustrates a TAM in contact with a TRPC, with both cell types containing lipids. This observation suggests a possible interaction and metabolic exchange between TAMs and TRPCs. Based on these results, we propose a model in which the GBM microenvironment comprises distinct micro-niches specifically driven by TSCs and TRPCs. TSC areas are infiltrated by cells of lymphoid origin capable of tumor targeting and killing, while TRPC regions, distinguished by specific metabolic features such as increased lipid metabolism, recruit immunosuppressive TAMs. These TAMs not only shape immunosuppressive micro-niches but also provide metabolic support to TRPCs by mediating and facilitating lipid trafficking (***Fig. 3U***).

**Figure 3:**
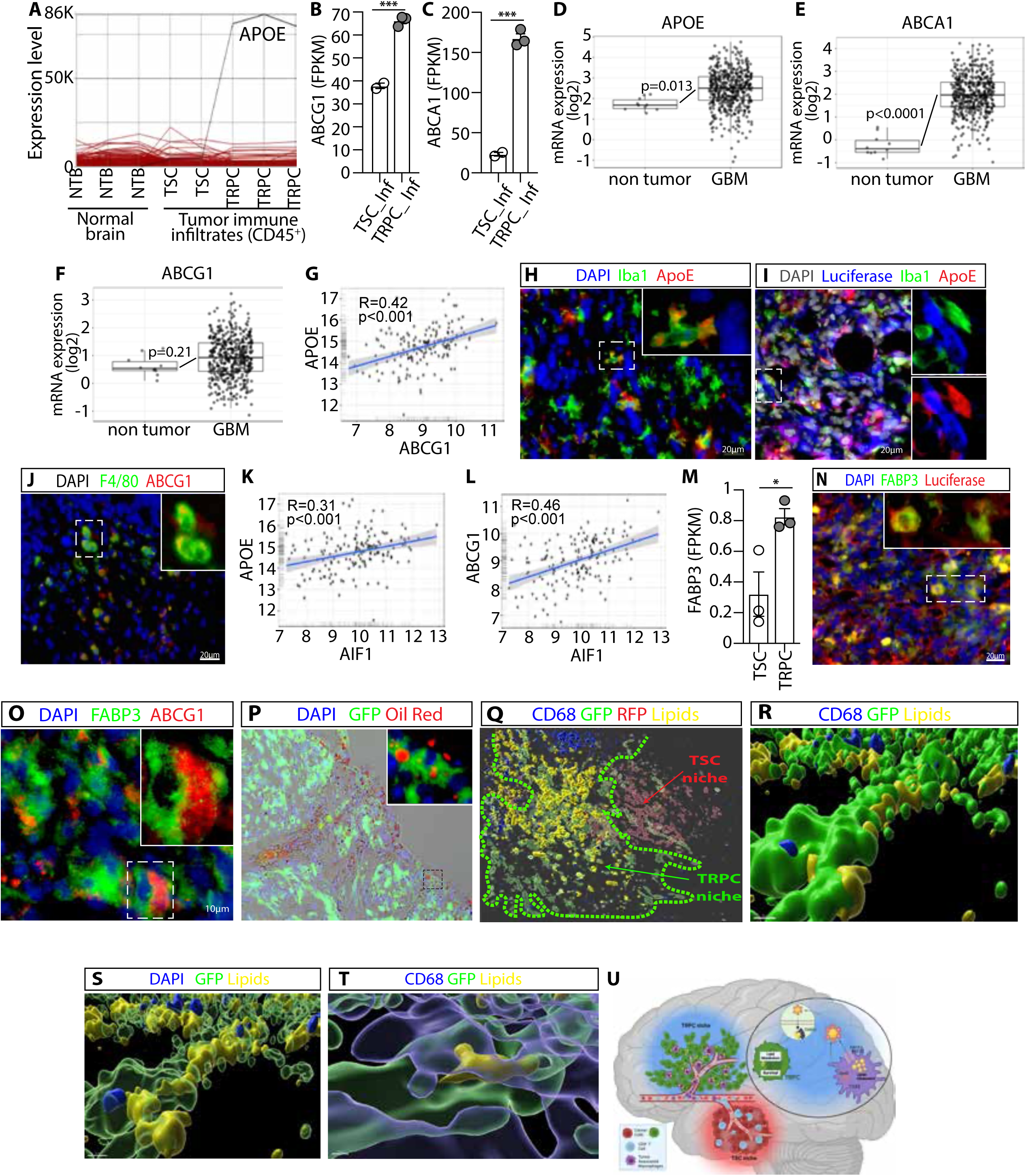
Metabolic Niches of TRPCs Are Defined by Lipid Accumulation and Immune Remodeling. **A-C)** Differential transcriptomic expression of (A) APOE, (B) ABCG1, (C) ABCG1 in TRPC ImmInfs; (Unpaired t-test) **D-F)** Transcriptomic expression of (D) APOE, (E) ABCA1, (F) ABCG1 in human GBM (hGBM). (Unpaired t-test) **G)** Pearson correlation analysis of APOE and ABCG1 transcript expression in hGBM **H-J)** Representative immunofluorescence staining of mTRPC tissue: (H) Nucleus, TAMs and APOE, (I) Nucleus, Tumor cells, Macrophages and APOE, (J) Nucleus, F4/80, ABCG1 (n=3) **K-L)** Pearson correlation analysis between: (K) APOE and AIF1, (L) ABCG1 and AIF1 **M)** FPKM values for FABP3 expression (Unpaired t-test) **N-P)** Representative immunofluorescence staining of mTRPC tissue: (N) Nucleus, Tumor cells, FABP3, (O) Nucleus, FABP3, ABCG1, (P) Nucleus, GFP+TRPCs, Red Oil (n=3) **Q-T)** 3D rendered images of mTME: (Q) GFP-TRPCs, RFP-TSCs, TAMs, Lipid Droplets, (R) Zoomed-in image of Q showing Lipids in TRPC region, (S) Transparent filter view of TRPCs containing lipids, (T) Higher magnification view of (R) showing spatial relationships among lipids, TAMs, and TRPCs (representative of 5 different brains analyzed) **U)** Proposed GBM TME model

### Spatially resolved TAM enrichment correlates with TRPC niches in human GBM

To validate our proposed model in the context of human disease, we analyzed GBM patient tissue samples to conduct detailed spatial mapping and characterization of the immune micro-niche within the TME. We performed spatial proteomic profiling based on hyperplexed sequential immunofluorescence using a 10-marker panel (***Fig. 4A*, *S4A***) ^56^. TRPC regions were defined from tissue samples of 5 different GBM patients based on the strong expression of FABP7 (***Fig. S4B)***, which we previously reported has being strongly upregulated in this cell lineage in human GBM (**EMBO**). Conversely, low FABP7 positivity was used to identify TSC areas. Immune contexture was characterized based on the presence of T lymphocytes (CD31-/CD3+), APCs (CD31-/HLA-DR+), cytotoxic T cells (CD31-/CD68-/CD4-/CD8+), T helper cells (CD31-/CD68-/CD4+), MDSCs (CD31-/CD68+/CD11b+/HLA-DR-), and TAMs (CD31-/CD68+)(***Fig. S4C***). TAMs were further subclassified based on the expression of FABP7 to identify those specialized in lipid homeostasis and metabolism, referred to as lipid-specialized TAMs (LS-TAM) (***Fig. S4D-I***) ^57–61^. A total of 39 ROIs were investigated to assess the spatial and phenotypic differences between the two tumor regions (***Fig. S4B)***. TSC regions demonstrated a higher presence of APCs and lymphocytes, particularly cytotoxic T cells and Th cells, indicative of a more immunogenic “hot” microenvironment (***Fig. 4B-E***). In contrast, TRPC-enriched regions exhibited a significant enrichment in LS-TAMs and a trending increase in MDSCs, suggesting a more immunosuppressive “cold” niche characterized by lipid homeostasis (***Fig. 4F-G***). Confocal 3D imaging revealed high density of immunosuppressive M2-like TAMs (CD163^+^) within TRPC-enriched regions, with proximity and frequent cell-cell contact between these TAMs and TRPC cells, suggesting a direct role in immune modulation and tumor progression (***Fig. 4H, I***). Additionally, single-cell neighborhood analysis, was performed on the same five GBM patient tissue samples using the analytical tool Single-cell Spatial Neighborhood Analysis and Quantification (SNAQ^TM^) to quantitatively assess LS-TAMs distribution within TRPC and TSC regions of human GBM ^14^. Staining for DAPI (nuclei), FABP7^high^ (TRPCs), FABP7^low^ (TSCs), and CD163 (TAMs), along with pseudo-coloring following cell identification and segmentation, are shown (***S4J-L***). TRPC neighborhoods were significantly enriched in M2-TAMs (***Fig. 4J***) with a significantly reduced distance to the closest TAM or 10 closest TAMs compared to TSCs (***Fig. 4K*, *S4M***). These observations indicate a distinct immune infiltration pattern, marked by the preferential accumulation of M2-TAMs within TRPC-enriched regions which aligns with our observations in the murine model, further supporting the role of TRPCs in shaping an immunosuppressive TME.

**Figure 4:**
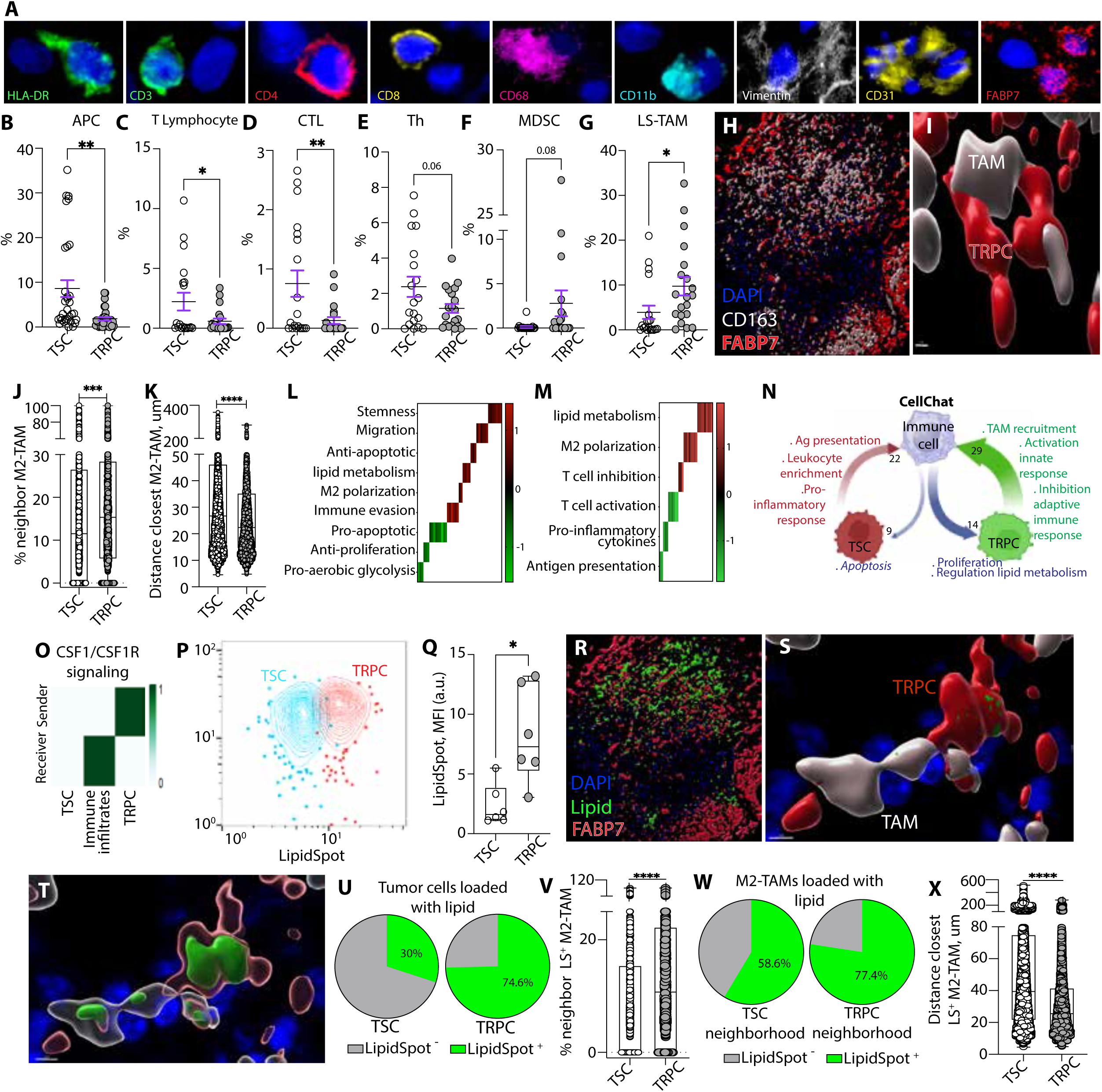
Spatial multi-omics reveals lipid-specialized TAM infiltration in TRPC niches of hGBM. **A)** Representative images of cells expressing markers: HLA-DR, CD3, CD4, CD8, CD68, CD11b, Vimentin, CD31, FABP7 (n=5) **B-G)** Percentage of immune cell types in hTRPC/hTSC ROIs: B) APC, C) T Lymphocytes, D) CTL, E) Th, F) MDSC, G) LS-TAM (n=5; Unpaired t-test) **H-I)** Representative 3D rendered image of hGBM TME (n=5). (H) Nucleus, Macrophage and TRPCs, (I) Magnified image **J-K)** Neighborhood analysis: (J) Percentage of M2-TAMs, (K) Distance of closest M2-TAMs to GBM cells; (Unpaired t-test) **L-M)** Heatmap of DEGs defining functions: (L) in immune cells, (M) in tumor cells (Log2Fc[TRPC/TSC]) **N)** Schematic of ligand-receptor bidirectional interactions between ImmInfs and hTRPCs and hTSCs (n=4) **O)** Quantitative sender-receiver communication heatmap for CSF1/CSF1R axis (n=4) **P)** Flowcytometric contour plot showing lipid spot intensities in hTRPCs and hTSCs (n=6) **Q)** Bar diagram quantification of P; (n=6; Paired t-test) **R-T)** Representative 3D image of hGBM TME (n=5): (R) Nucleus, Lipid droplets and TRPCs, (S) Magnified image of R, (T) Transparent filter of S **U-X)** Corresponding neighborhood analysis (n=5) of panel 4R-T. (U) Mean percentage of lipid loaded tumor cells, (V) percentage of neighbor loaded M2-TAMs; Unpaired t-test. (W) Mean percentage of lipid loaded M2-macrophages cells, (X) Distance of closest lipid loaded M2-macrophages in hTME; (Unpaired t-test)

### Transcriptional regulation and intercellular signaling define TRPC immune landscape in human GBM

To explore the molecular drivers of cellular heterogeneity and gain insights into cellular composition and activity, we performed spatial transcriptomic phenotyping. The GeoMx Whole Transcriptome Atlas (WTA) was utilized to capture the full spectrum of transcripts from 92 ROIs of the same five human GBM biopsy samples (***Fig. S5A***). Through DEG analysis, we identified differentially regulated functions within TRPC specific regions (***Table S6***). TRPC segments were enriched in genes related to stemness, cell migration, immune evasion, M2-like macrophage polarization, and lipid metabolism while pro-aerobic glycolysis genes were downregulated, aligning with anticipated metabolic adaptations (***Fig. 4L*, *Supp. table S6***) ^11^. The transcriptomic profiles of TRPC-enriched immune segments showed upregulation of genes linked to the M2 macrophage phenotype, T cell inhibition, and lipid metabolism, with downregulation of genes related to T cell function, pro-inflammatory cytokines, and antigen presentation (***Fig. 4M***). Together, these results strongly corroborate our previous studies from our murine model, linking TRPC with a cancer stem cell phenotype characterized by specific metabolic traits, including upregulated lipid metabolism ^11,12^, and associated with specific immune microniches. This study identified transcriptional programs governing immune composition and diversity, metabolic specificities, and cell-cell interactions, providing a comprehensive framework for understanding tumor-immune dynamics and potential therapeutic targets in human GBM. Importantly, these findings highlight the spatially resolved nature of these characteristics.

Next, we deciphered the differential cellular communication network and enriched biological interactions particularly between TRPCs and their surrounding immune infiltrates, utilizing the CellChat platform. This tool deconvolutes complex intercellular communications from scRNA-seq data by analyzing ligand-receptor, cofactor, and membrane-bound receptor expression at single-cell resolution ^13,57^. We analyzed sample data from four GBM patients ^62^ where TSCs and TRPCs were identified and classified as previously described ^13^. TRPCs and immune cells engage in 43 unique ligand–receptor interactions, with 29 ligands expressed by TRPCs and 14 by immune cells (***Fig. 4N, S5B, C*, *Tables S7 and S8***). In contrast, TSCs share only 31 ligand– receptor pairs with immune cells, comprising 22 ligands from TSCs and 9 from immune cells. Using the STRING database ^63^ (https://string-db.org), we found that one of the most enriched communication pathways between TRPCs and immune cells is the MCSF signaling (GO:0038145), which supports the differentiation and survival of macrophages. Our results show that the CSF1/CSF1R axis is key in TRPC-immune cell communication, with TRPCs as the primary source of CSF1 and immune cells as the main recipients (***Fig. 4O*, *S5B***). CSF1 alone drives 100% of CSF1R activity, with no contribution from IL-34 ^64–67^ (***Fig. S5D***). Additional key signaling pathways activated by TRPC-mediated signals to immune cells include the positive regulation of microglial cell migration (GO:1904141), positive regulation of macrophage migration (GO:1905523), and macrophage chemotaxis (GO:0010759). Moreover, TRPCs contribute to immune suppression by upregulating pathways involved in the negative regulation of T cell-mediated cytotoxicity (GO:0001915) and dendritic cell differentiation (GO:2001199). Thus, by promoting macrophage migration and recruitment while concurrently suppressing adaptive immunity, TRPCs contribute to tumor progression and immune evasion, hallmarks of a “cold” immune microenvironment. These results strongly align with our animal model data and spatial proteomics and transcriptomics analyses in human GBM, further reinforcing the central role of TRPCs in tumor-immune interactions. Conversely, the signals transmitted by TSCs to immune cells were distinct and primarily associated with promoting a pro-inflammatory, “hot” immune microenvironment. These included pathways involved in leukocyte migration in response to inflammation (GO:0002523), leukocyte chemotaxis in inflammatory responses (GO:0002232), leukocyte aggregation (GO:0070486), cellular response to interferon-alpha (GO:0035457), antigen processing and presentation of endogenous peptides (GO:0002476), positive regulation of NK cell differentiation (GO:0032825), and enhanced NK cell cytokine production (GO:0002729). Notably, TSCs also upregulated pathways involved in hematopoietic stem cell (HSC) migration to the bone marrow (GO:0097241). Increased intratumoral presence of HSCs has been linked to enhanced anti-tumor immunity ^19^. These findings suggest that TSCs orchestrate a robust pro-inflammatory response, fostering an immune microenvironment that strengthens tumor surveillance and counters immune evasion. In terms of signaling from immune cells to tumor cells, TSCs primarily receive pro-inflammatory cues that activate the extrinsic apoptotic pathways via the death domain receptor (GO:0008625). However, immune cells primarily communicate with TRPCs through signals that promote cell proliferation (GO:0038134, GO:0060252, GO:0061900) and, notably, regulate lipid metabolism (GO:0060559, GO:0030730). The latter enhances the tumor cells’ ability to uptake and sequester lipids (GO:0030730), further supporting the findings observed in murine models.

### Spatial co-localization between TRPC niches, lipids, and lipid specialized TAMs in human GBM

Based on these findings, human TRPCs, like those in murine models, are expected to contain high lipid levels and be spatially associated with lipid-enriched regions characterized by a dense TAM network. To confirm and functionally validate these associations, we quantified lipid content in TRPCs compared to TSCs within tumors isolated from six different GBM patients. Using chip cytometry, we extracted quantitative data from IHC images (***Fig. S5E).*** Analysis revealed significantly higher lipid content in TRPCs compared to TSCs (***Fig. 4P***, ***Q, S5F-I***), further supporting in situ their distinct metabolic profile. Building on our findings, we sought to further illustrate the differential spatial distribution of lipids within the human GBM microenvironment. To achieve this, we employed 3D reconstructions to map lipid droplets distribution and further confirmed preferential accumulation in TRPC regions and increased presence of M2-like TAMs (***Fig. 4H, R, S5J***). Zoomed-in images revealed lipid-containing TAMs and lipid-containing TRPCs in intertwined connection suggesting a spatially organized, metabolically driven interaction between these cell types (***Fig. 4S, T*, *Video S3***). SNAQ™ analysis revealed 74.6% of TRPCs contained lipids, compared to 30% in TSCs (***Fig. 4U***). Within a 55um radius of TRPCs, 14.9% of cells are lipid-loaded M2-like TAMs, compared to only 9.8% in TSC regions (***Fig. 4V***). Notably, 77.4% of TAMs in TRPC neighborhood are lipid-loaded, compared to 58.6% in TSC areas (***Fig. 4W***). TRPCs also exhibited shorter distances to the nearest lipid-containing M2-like TAMs (***Fig. 4X***).

Therefore, using spatial proteomics, transcriptomics, and advanced profiling, we characterized TRPC micro-niches in the human GBM TME. Altogether, these data support that TRPC micro-niches in human GBM, similar to those observed in murine models, are comprised of lipid-rich tumor cells that are heavily infiltrated by immunosuppressive TAMs, creating a ’cold’ immune microenvironment that promotes tumor progression, facilitates immune evasion, and provides essential metabolic support.

### Macrophage-mediated lipid transfer provides metabolic support that fuels TRPCs

Given that TRPC regions of GBM are characterized by high lipid content in both tumor cells and infiltrating lipid-specialized M2-like TAMs, we hypothesize that lipid trafficking from TAMs to tumor cells supports TRPC metabolism, promoting tumor growth, survival, and progression. Our next objective was to provide evidence of lipid transfer in the human GBM TME. Using IMARIS colocalization and surface tools with 3D-rendered imaging, we identified colocalization of lipids with TAM-TRPC contact points, providing a snapshot of potential lipid exchange (***Fig. 5A-C***). Although time-lapse imaging is needed to measure transfer kinetics, these results showed continuous lipid localization from TAM cytoplasm to TAM/TRPC membranes and into TRPCs, suggesting lipid transfer at points of contact, though trafficking directionality remains unclear.

**Figure 5:**
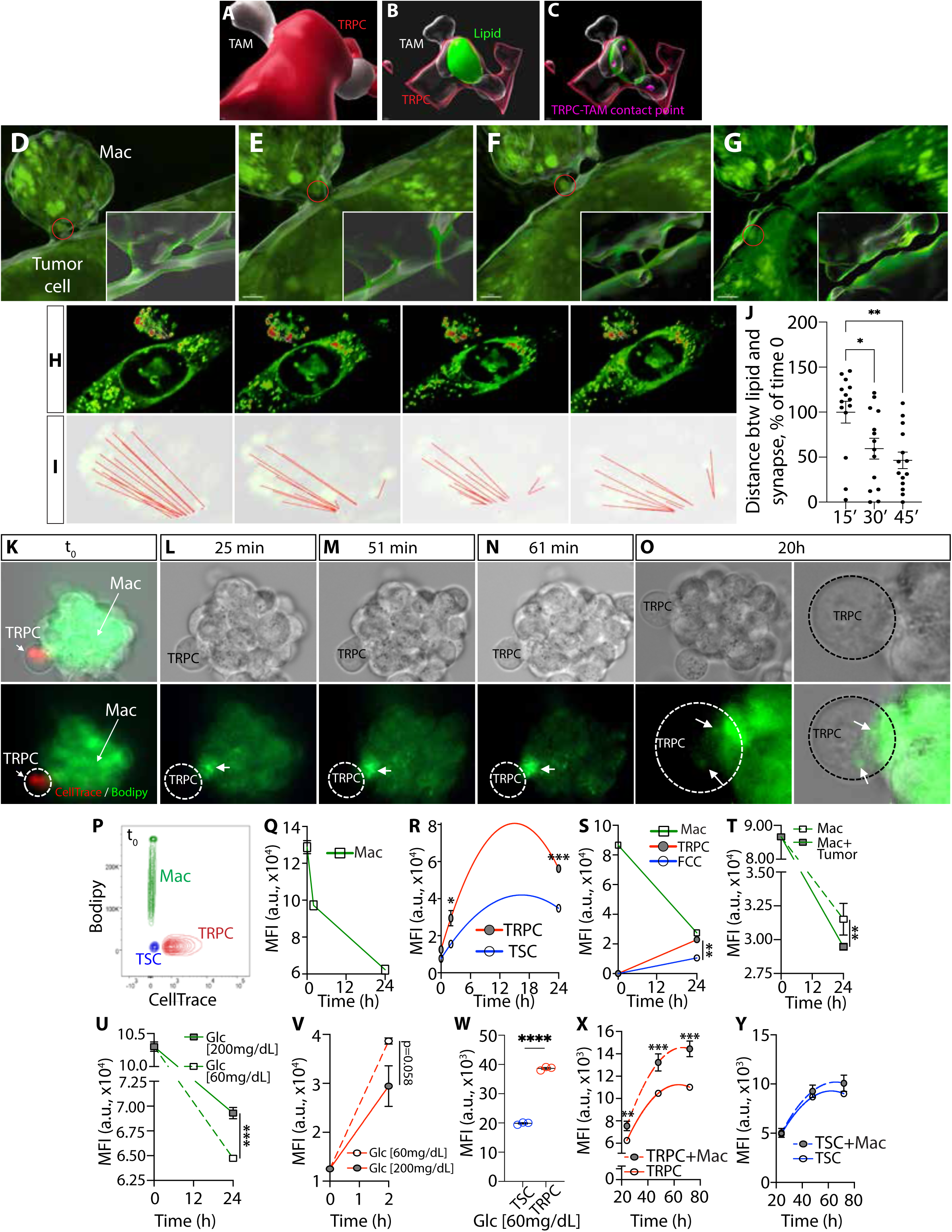
Supporting TRPC growth through macrophage lipid transfer. **A-C)** Representative 3D rendered images in hTME: (A) FABP7^+^TRPCs contact with CD163^+^TAM. Transparent filter showing (B) LipidSpot^+^Lipids, and (C) TRPC–TAM interfaces as potential sites of lipid transfer; (n=5) **D-G)** Representative 3D label free refractive index based holotomographic images showing stages of KLuc and IC21 contact with lipid transfer. Insets: Zoomed-in contact points (n=3) **H-J)** Representative time-lapse holotomographic pseudocolored-images at 24h, 24h+15mins, 24h+30mins, 24h+45mins showing: (H) lipid movement, (I) Red lines showing distance of lipid from the KLuc-IC21 contact, (J) Dot plot of 5I; (n=3; Unpaired t-test) **K-O)** Time lapse imaging of lipid transfer from IC21 to mTRPC at times: (K) 0, (L) 25 mins, (M) 51 mins, (N) 61 mins, (O) 20h **P-U)** Flow cytometry of lipid trafficking in 24 hours: (P) Contour plot showing lipid content at time= 0, (Q) Lipid efflux from THP-1s, (R) Lipid influx to human TRPCs and TSCs, (S) Overall lipid efflux from IC21s to murine TRPCs and TSCs, (T) Lipid efflux rate from IC-21 cells with or without tumor cells, (U) Lipid efflux from THP-1 cells under different glucose conditions in presence of tumor cells; (n=3, 3 Unpaired t-test) **V)** Lipid uptake rate in hTRPCs in different glucose conditions **W)** Lipid uptake under glucose-limited environment in hTRPCs and hTSCs (n=3,3; Unpaired t-test) **X, Y)** Cell viability of (X) hTRPCs, and (Y) hTSCs co-cultured with THP-1 macrophages under low glucose conditions (n=4, 4; Unpaired t-test)

To address this, we utilized time-lapse holotomography capturing real-time imaging of endogenous lipids. IC21 murine macrophages were co-cultured with KR158 cells and time-lapse acquisition revealed a multistep process of endogenous lipid transfer from a macrophage to a tumor cell (***Fig. 5D-G*, *S6A-B***). The initial image captures the latching phase, where both cell types establish contact. By 15 minutes, a bridge forms between the macrophage and the tumor cell, facilitating lipid exchange. At 30 minutes, the lipid droplet transfer is actively underway, progressively moving from the macrophage into the tumor cell. By 45 minutes, detachment occurs, with the lipid droplet fully internalized within the tumor cell, signifying successful intercellular lipid transfer. Interestingly, the relative distribution of lipids within the macrophage shifted over time, with lipids mobilizing toward the cell-cell contact site before transfer. This directional mobilization of the lipid network resulted in a significant decreased distance between lipids and the point of contact (***Fig. 5H-J*, *S6A***). Additionally, a cellular protrusion was noted specifically extending from the macrophage latching onto the tumor cell (***Fig. 5D-G*)**. Together, these findings suggest an asymmetric lipid flux directed from macrophage to tumor.

To characterize further this phenomenon, we co-cultured C16-BODIPY-labeled THP-1 monocytes with human TRPCs and monitored exogenous lipid transfer via time-lapse imaging (***Fig. 5K***). Within 25 minutes, lipid redistribution within the THP-1 cell network toward TRPCs was also observed, intensifying over time (***Fig. 5K-O***). After 20 hours, lipid transfer occurred, with agglomerates forming in TRPCs by 44 hours (***Fig. 5O*, *S6C-D***). After 48 hours, multiphoton microscopy offered a complementary comprehensive perspective of lipid distribution and transfer, reinforcing directional lipid trafficking from THP-1 cells to TRPCs (***Video S4***), with similar results in murine macrophages and glioma cells (***Fig. S6E-F***). Time-course flow cytometric studies were conducted to quantify the differential lipid transfer efficiency between THP-1 cells with human TRPCs and TSCs (***Fig. 5P-R*, *S6G***). At t=0, C16-BODIPY fluorescence was exclusively detected in THP-1 cells. Over time, lipid efflux from THP-1 cells (***Fig. 5Q***), resulted in progressive lipid acquisition by TRPCs and TSCs (***Fig. 5R***). Notably, TRPCs exhibited a significantly higher lipid uptake rate (***Fig. 5R*, *S6H-I***). Similar results were observed in the mouse system after co-culturing IC-21 macrophage cells with KR158-derived TRPCs and TSCs (**Fig. 5S, S6J-K**). Next, we observed a significantly higher lipid efflux rate when murine IC-21 macrophages were co-cultured with tumor cells compared to macrophages cultured alone over a 24-hour period (***Fig. 5T***). This indicates that tumor cells actively induce intercellular lipid transfer.

To explore lipid trafficking under metabolic pressures reflective of the TME, we examined the impact of low-glucose conditions (60 mg/dL) ^68^, on lipid trafficking dynamics. In presence of human GBM cells, low-glucose conditions significantly stimulated lipid efflux from THP-1 macrophages compared to physiological glucose levels (200 mg/dL) (***Fig. 5U***). Notably, macrophages cultured alone under similar conditions exhibited no significant change in lipid efflux (***Fig. S6L***), suggesting that lipid mobilization is not solely driven by glucose availability but rather is a regulated metabolic adaptation influenced by cell interactions in response to the metabolic status of their surrounding microenvironment. To rule out if metabolic stress-induced cytotoxicity could affect macrophage lipid efflux or induce macrophage death releasing lipids, we performed CyQUANT™ Cell Assay to assess macrophage viability at 48 and 72 hours. No cytotoxic effects were observed (***Fig. S6M-N***), suggesting that this process of lipid transfer reflect a broader metabolic strategy by tumor cells to adapt to the TME by acquiring lipids to support growth, survival, and possibly immune evasion. Importantly, when co-cultured with THP-1 macrophages, TRPCs increase their lipid uptake in response to low-glucose concentration (***Fig. 5V***), incorporating significantly greater level of lipids compared to TSCs (***Fig. 5W***). This suggests that TRPCs can adapt to metabolic stress by upregulating lipid trafficking and acquisition from macrophages to meet their metabolic demands. Notably, under low-glucose conditions the presence of macrophages significantly increased the number of TRPCs over time (***Fig. 5X***) while macrophage co-culture had no impact on TSC survival (***Fig. 5Y***), suggesting a selective advantage conferred to TRPCs through macrophage-mediated metabolic support.

### FABP3-mediated lipid homeostasis regulates the tumor metabolic and immune environment modulating tumor progression

Our next objective was to establish the functional significance of this metabolic homeostasis in driving tumor growth by disrupting this key pro-tumorigenic mechanism. Our previous studies showed the central role of FABPs, more particularly FABP7 and FABP3, in regulating lipid metabolism and uptake in GBM ^11^. In this study we assessed *in vivo*, the effect of decreasing lipid uptake via FABP3 inhibition by administering a pharmacological inhibitor intracranially into tumor-bearing mice ^11^. Immunohistochemistry, combined with 3D rendering and lipid surface measurement, revealed that the treatment led to a significant reduction in lipid quantity, a more confined distribution of lipids within the TME accompanied by a decrease in intratumoral TAM infiltration (***Fig. 6A-E*, *S7A***). Single-cell spatial quantitative cellular neighborhood analysis using SNAQ^TM^ ^69^ showed a 2.5-fold reduction in TAM presence and increased distance to the nearest TAMs (50.76 µm vs. 36.96 µm in controls) in response to treatment [***Fig. 6F, G, S7B-G***]). The distance to lipid-loaded TAMs from tumor cells also increased 1.7 times with inhibitor treatment (***Fig. 6H***). Importantly, these changes in the TME correlated with delayed tumor growth and increased survival (***Fig. 6I, J*)**. Our study identified intratumoral lipid homeostasis, driven by TRPCs and TAMs, as a key factor in GBM growth. Targeting this regulatory network to restrict lipid availability may disrupt disease progression and improve patient outcomes.

**Figure 6:**
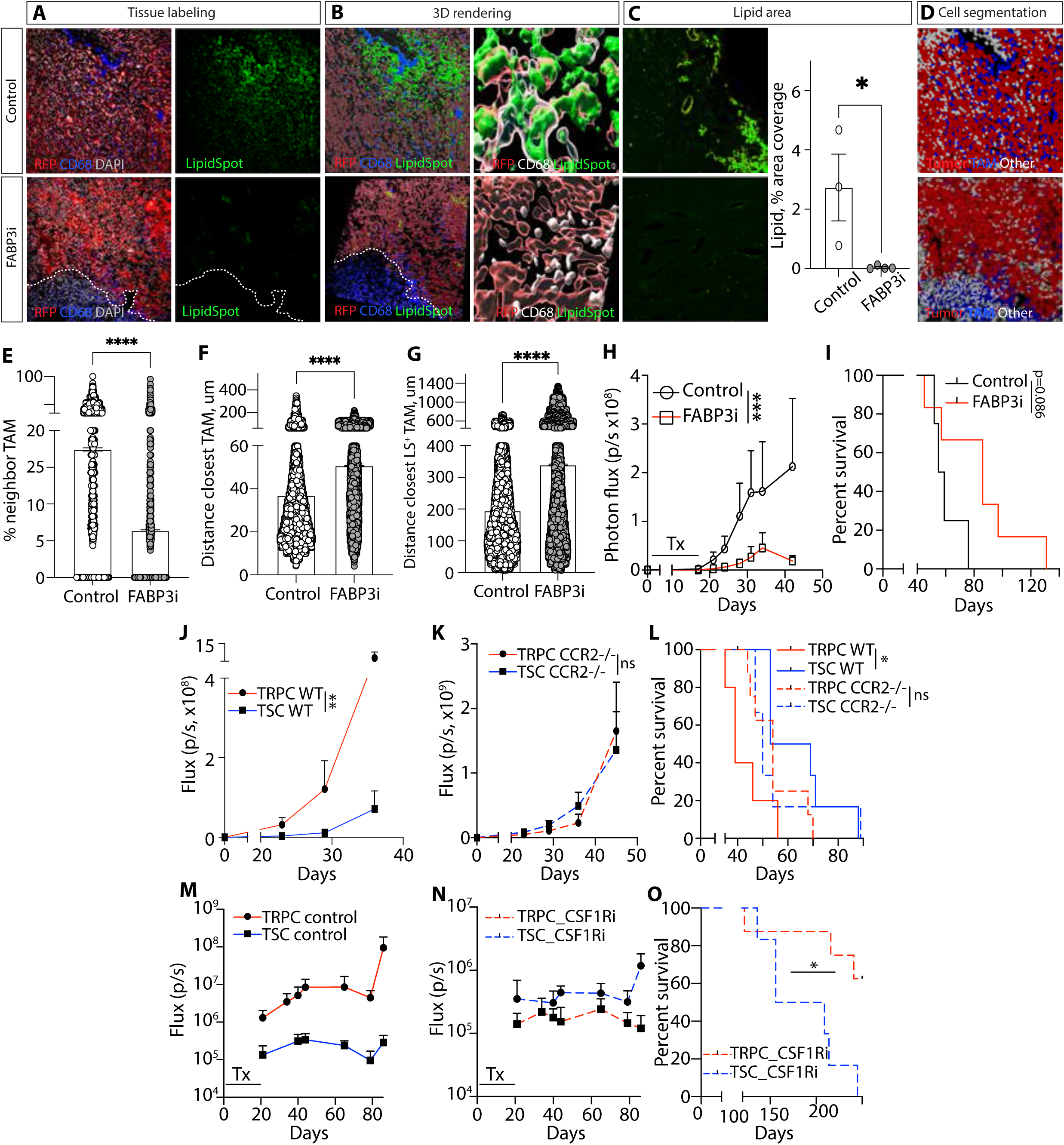
Disrupting TRPC-TAM metabolic homeostasis impairs tumor growth. **A-D)** Representative IHC images taken under standard imaging conditions of mTME upon SB-FI-26 treatment (A) Tissue labeling with RPF^+^ tumor cells, macrophages, nuclear staining and Lipids, (B) 3D rendered TME (n=3 for Control and 4 for FABP3i treated), (C) Representative tissue images showing lipids identified by uniform thresholding using Fiji/ImageJ and its corresponding bar-diagram showing percent lipid coverage, (D) Cell segmentation. (Dotted line demarcates the TME from the non-TME region) **E-G)** Neighborhood analysis showing (E) percentage of TAM, (F) distance of closest TAM, and (G) Distance of closest lipid loaded TAM from tumor cells (n=3, 4, Unpaired t-test) **H-I)** (H) *In vivo* monitoring of intracranial tumor growth (n=5, 7), (I) Kaplan-Meier survival curves in Control vs SB-FI-26 treated mice (n=4, 6; Log-rank test) **J-K)** *In vivo* monitoring of intracranial tumor growth of TRPC and TSC in (J) WT (n=5,6) and in (K) CCR2 KO mice (n=9, 6; two-way ANOVA). L) Kaplan-Meier survival curves (WT n=5, 6; KO n= 9, 6; Log-rank test) **M-N)** *In vivo* monitoring of intracranial tumor growth of TRPC and TSC in (M) Control (n=9, 7) and (N) CSF1Ri treated mice (n=7,5; two-way ANOVA). **O)** Kaplan-Meier survival curve (n=8, 6; Log-rank test)

Next, we aimed to demonstrate *in vivo* that reduced intratumoral TAM presence specifically affects TRPCs. By targeting pathways that recruit and sustain macrophages (***Fig. 2T***), we found that CCR2-KO mice had significantly reduced MDSC and macrophage infiltration in the glioma TME ^18,70–74^. Using CCR2-KO mice, we assessed tumor progression in TRPC-and TSC-derived tumors under conditions of diminished macrophage infiltration. TRPCs formed more aggressive tumors in wild-type mice (***Fig. 6K***), but in CCR2-KO mice, the tumor growth advantage was negated and survival were similar between both groups (***Fig. 6L, M***).To further investigate the role of TAMs in TRPC-driven tumor progression, we used the InVivoMAb anti-mouse CSF1R antibody, for intratumoral TAM depletion ^38,37^; ^75^; ^76^. Intracranial blocking of CSF1R led to a significant reduction in tumor growth and extended survival specifically in the TRPC group (***Fig. 6N-P***). Together, these findings highlight the critical interplay specifically between TRPCs and immunosuppressive TAMs, demonstrating its role in tumor progression. Moreover, these results reveal that disrupting key cellular elements, which we demonstrated regulate lipid accumulation within TRPCs, through targeted immunomodulation results in significant therapeutic benefits.

### TRPC transcriptional signature correlating with specific immune profiles, predicts disease evolution and response to hypolipidemic drugs

To further assess the translational potential of our findings, we investigated the prognostic value of the TRPC-specific transcriptomic profile, including its related immune response signature linked to their lipid-specialized and TAM-rich environment identified in our murine model (***Fig. 2Q***). GSEA analysis of TRPCs and TSCs from nine GBM patients revealed that 35 of 54 immune response genes were significantly overexpressed in human GBM TRPCs (***Fig. 7A*, *Table S9, S10***). Furthermore, stratifying GBM patients from the TCGA dataset based on the expression of these genes demonstrated a strong association between high expression levels and poorer prognosis (***Fig. 7B***). Further supporting the results from our murine experiments, we also observed a significant positive correlation between CCL2 and CCL7 expression and markers of TAMs, including AIF1, CD163, MRC1, and CCR2, in human GBM (***Fig. S8A***). Moreover, elevated expression of these markers was associated with shorter survival times in GBM patients (***Fig. S8B***), suggesting that increased CCL2/CCL7 levels may contribute to poor prognosis by promoting TAM recruitment and maintenance. These findings highlight the clinical significance of the identified immune signature and its associated transcriptional correlations. The expression of the TRPC signature and its related immune response signature could serve as potential diagnostic or classification tools.

**Figure 7:**
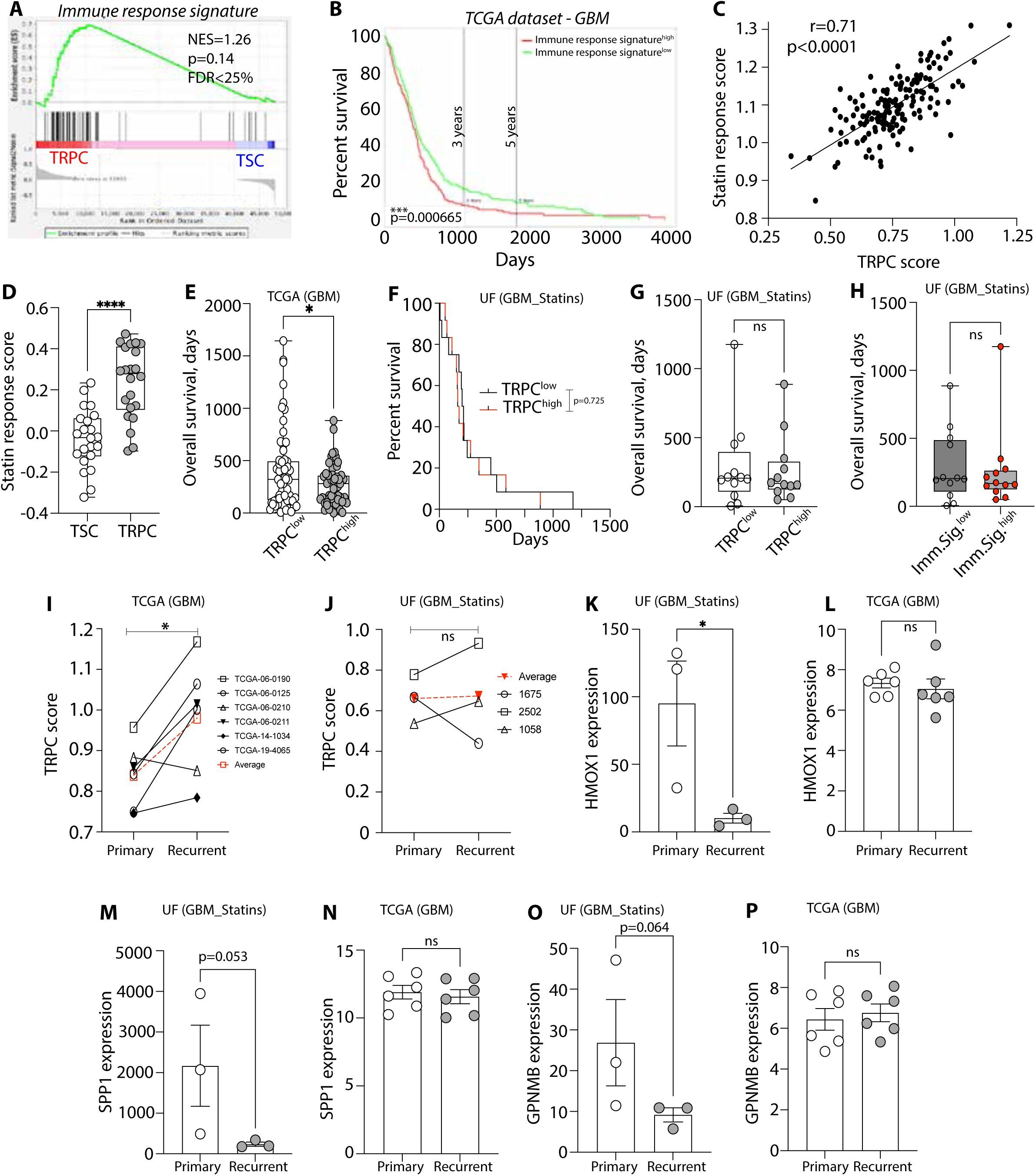
GBM patients with high expression levels of the TRPC transcriptional signature respond to hypolipidemic drugs. **A)** GSEA of Immune response signature in TRPC/TSCs from 9 GBM patients **B)** Percent survival of GBM patients (TCGA dataset) stratified based on immune response signature (median) **C)** Correlation between TRPC and Statin response score in hGBM (TCGA database) **D)** Statin response score in TSCs and TRPCs (scRNA seq data, 4 GBM patients) **E)** OS of GBM patients stratified based on TRPC signature (TCGA dataset) **F)** Kaplan-Meier survival curve showing survival for statin treated UF patients with high/low TRPC score (n=12, 12; Log-rank test) **G)** OS of 5F (n=12,12; Unpaired t-test) **H)** OS for statin-treated UF patients with high/low immune signature (n=12, 12; Unpaired t-test) **I-J)** TRPC scores of paired primary vs recurrent GBM patients from (**I)** TCGA database (n=4) and (**J)** UF patients (n=3; Paired t-test) **K, P)** Expression levels (TPM) of HMOX1 (K, L), SPP1 (M, N), and GPNMB (O, P) were compared between paired primary and recurrent GBM tumors from statin-treated UF patients cohort (panels K, M, O; n = 3) and from the TCGA dataset (panels L, N, P; n = 6, paired t-test)

Our findings suggest that GBM patients with a TRPC-associated transcriptomic signature may benefit from lipid metabolism-targeting treatments, like statins. Raghu *et. al.* developed a gene expression signature to differentiate statin-sensitive and resistant cells ^77^. Utilizing this statin-sensitive gene signature of 26 genes (***Table S11***), we found a strong positive correlation between the TRPC score and statin sensitivity in GBM patients from the TCGA database (***Fig. 7C***). Additionally, single-cell RNA sequencing of tumor tissue from four GBM patients showed higher expression of this statin sensitivity signature in TRPCs compared to TSCs (***Fig. 7D***), supporting the hypothesis that GBM patients with high TRPC scores may indicate greater statin response. To test this hypothesis, we conducted a retrospective study to analyze the potential correlation between TRPC scores and survival in GBM patients who were also treated with hypolipidemic drugs. We evaluated 24 patients with survival information and access to resected specimen, for RNA sequencing and TRPC signature-based stratification. Patients in the TCGA dataset characterized by a high TRPC signature exhibited significantly reduced median survival ^13^ and overall survival (***Fig. 7E***), consistent with a more aggressive tumor phenotype. In contrast, our retrospective analysis revealed that among patients receiving statin treatment, this survival difference was no longer observed. Specifically, TRPC^high^ and TRPC^low^ patients showed comparable survival outcomes in the statin-treated group (***Fig. 7F, G***). These results suggest that statin use may mitigate the poor prognosis typically associated with high TRPC expression. A similar phenomenon was observed when patients from the statins-treated cohort were stratified by the immune response signature, the survival difference between TRPC^high^ and TRPC^low^ tumors was no longer apparent, indicating that statins use may mitigate the aggressive progression associated with TRPC^high^ tumors (***Fig. 7H***). Moreover, in statin-treated patients, no correlation was observed between the TRPC score and survival (***Fig. S8C***), which is shown to be significantly and negatively correlated in TCGA GBM patients (***Fig. S8D***). This suggests that statin treatment may benefit patients with high TRPC expression. The TRPC transcriptomic characteristics have also clinical relevance in the context of disease recurrence. Analysis of the TCGA database revealed that recurrent GBM overexpressed this TRPC signature compared to their matched primary tissues (***Fig. 7I***). However, this enrichment is not observed in GBM patients treated with statins (***Fig. 7J***). This suggests that statin treatment can impact the TRPC cell population, potentially preventing the upregulation of TRPC-associated pathways and tumor phenotypes linked to therapy resistance, aggressiveness, and recurrence. These observations indicate that disrupting the tumor metabolic homeostasis via statins treatment may alter the molecular landscape of GBM, potentially affecting tumor aggressiveness and recurrence patterns. This effect is specifically observed in GBM patients with a high TRPC score, suggesting that this signature could be used to identify patients who are responsive to statin treatment. Interestingly, our retrospective analysis of statin treatment in GBM patients revealed that, upon tumor recurrence, key genes within the lipid-laden macrophages (LLMs) signature, which has been linked to resistance to immune checkpoint inhibitors, exhibited notable reductions (***Fig.7K-P***) ^78^. This reduction was not observed in recurrent GBM cases not selected for statin therapy (***Fig. 7L*)**. Significantly, this population of LLM exhibit overlapping phenotypic (e.g., immunosuppressive) and metabolic (e.g., lipid specialized) properties with the TAMs described in our study. These findings suggest that statins may serve as effective combinatorial or neoadjuvant agents to enhance sensitivity to immune checkpoint blockade (ICB), potentially by disrupting the metabolic crosstalk between TRPCs and TAMs through depletion of metabolically supportive TAMs. This aligns with recent work by Kloosterman and colleagues, who demonstrated that LLM burden can serve as a predictive biomarker for ICB response, and that decreasing lipid specialized TAMs correlates with improved therapeutic outcomes.

Collectively, these findings establish a compelling rationale for prospective clinical investigations assessing lipid-lowering agents, either as monotherapies or in combination with immune checkpoint inhibitors (ICI). Future GBM trial designs should incorporate the distinct transcriptional signatures identified in this study to enable biomarker-driven patient stratification and therapeutic response assessment.

## DISCUSSION

Our study leverages a systems biology framework integrated with spatial multi-omics and deep capture network approaches to provide a comprehensive view of the key regulators underlying fundamental mechanisms of GBM. We identified a critical metabolic crosstalk between peripherally recruited, metabolically distinct immunosuppressive TAMs and a specific tumor cell lineage linked to resistance to standard-of-care therapies and tumor recurrence (i.e., TRPCs) ^11,12^. Our findings suggest that TRPCs metabolically hijack these recruited immune cells to fulfill their pro-tumorigenic functions via facilitating lipid transfer and acquisition. Going beyond a model-based approach, we validated our observations in treatment-naïve human GBM tissue systems and demonstrated the clinical relevance of our results through the interrogation of multiple patient databases and conduction a retrospective clinical study linking TRPCs with potential response to lipid metabolism targeting therapeutic strategies.

The metabolic functions of macrophages, including lipid recycling, transport, and transfer, are well documented ^78–84^. A recent study elegantly demonstrated that, in the context of radiotherapy, TAMs can deliver cholesterol recycled from degraded myelin sheaths to GBM cells ^78^. While this mechanism appears to be specific to radiation-induced myelin breakdown, our findings suggest that GBM cells, particularly TRPCs, can metabolically co-opt TAMs even in the absence of therapy. This supports the idea that lipid-based metabolic crosstalk between GBM cells and immune cells represents a fundamental mechanism of tumor support and immune evasion, independent of treatment. Moreover, this interplay may extend beyond myelin recycling, potentially involving lipids derived from apoptotic or necrotic cells, or supplied through lipid-specialized brain-resident cells such as astrocytes or microglia, which may feed lipids to TAMs that, in turn, fuel tumor cells.

Our findings reveal that lipid metabolic dependencies, particularly those involving lipid availability, trafficking, and uptake, are critical players of tumor progression driven by TRPCs. We identified a distinct TRPC-associated metabolic gene signature that is strongly correlated with aggressive disease and poor prognosis in GBM patients. Importantly, in a retrospective clinical analysis, GBM patients with high TRPC metabolic scores who received statin therapy did not exhibit the reduced survival typically associated with this high-risk signature in untreated cohorts. These results suggest that the TRPC metabolic signature may serve as a clinically actionable biomarker to stratify patients and identify those most likely to benefit from statin therapy. Prior clinical studies investigating statins in GBM have largely yielded negative results, likely due to the absence of molecular stratification ^85^. Our study highlights the promise of leveraging TRPC-driven metabolic profiling to guide patient selection, improve therapeutic outcomes, and support the development of metabolism-based precision medicine strategies in GBM.

A recent study in lung cancer patients reported that concurrent statin therapy with ICB was associated with improved clinical outcomes ^86^ and other cancer types ^87–91^. While the precise mechanisms remain to be fully elucidated, the well-documented pleiotropic effects of statins, including their roles in modulating lipid metabolism and immune regulation, are likely contributors (^92, 93–96^. In our study, disruption of lipid homeostasis resulted in significant remodeling of the tumor immune microenvironment, marked by reduced infiltration of TAMs and delayed disease progression. Notably, statin treatment was associated with a decrease in the gene signature corresponding to lipid-laden TAMs, a population previously linked to resistance to ICB. These findings suggest that statins may counteract immune suppressive features of the tumor microenvironment, potentially enhancing the efficacy of immune checkpoint inhibitors. This supports the rationale for investigating combination therapies involving ICIs and statins, or broader strategies targeting lipid metabolism, to overcome immune resistance in GBM and other solid tumors.

## Limitations of the study

This study identifies specific pathways of TAM recruitment by TRPCs and demonstrates a functional role for TRPC-TAM lipid homeostasis in driving disease progression. While these findings uncover a critical axis of metabolic and immune interaction, the specific lipid species have not been fully characterized. Identifying the precise classes and sources of lipids accumulating in TAMs and transferred to TRPCs will be essential for delineating the molecular mechanisms underlying lipid-mediated immunosuppression and tumor progression. Future studies employing lipidomics, spatial metabolomics, and cell-type-specific tracer analysis will be important to resolve these details. Additionally, a more comprehensive exploration of lipid signaling functions, beyond metabolic and structural support, is needed to fully understand their roles in GBM malignancy. Finally, prospective, biomarker-guided clinical trials with statins alone or in combination with ICIs in patients stratified by TRPC metabolic signatures will be essential to validate the translational significance and therapeutic potential of this approach in the clinical setting.

## ACKNOWLEDGMENTS

We acknowledge Venkata Vijaya Karthik Narisetty, Mia Engelbart, and Ethan Hodge for their contributions in preparing cells for holotomography experiments, and Kaytora Long-James and Illena West for their excellent support in animal care and organ collection. We thank Benjamin Staley from the Integrated Data Repository Research Service at the University of Florida for assistance in retrieving clinical data for our retrospective study investigating GBM patients treated with lipid-lowering therapies. We are grateful to Daniel Silver, Florian Siebzehnrubl, and Brent Reynolds for their constructive and insightful comments and discussions, which greatly enriched this work. We are grateful to Andrea Doti from the University of Florida’s Interdisciplinary Center for Biotechnology Research for her guidance with flow cytometry experimental design and execution of cell sorting, and to Doug Smith for his support with microscopy and IVIS imaging. We also thank Bayli DiVita Dean from the Flores laboratory at the University of Florida for assistance with Luminex experiments. We thank Ann Dongtao Fu from the Molecular Pathology Core at the University of Florida for expert technical assistance with immunohistochemistry and GeoMx studies. We also would like to acknowledge Hinda Najem from the Heimberger laboratory at Northwestern University for valuable input on the SeqIF experiments using the COMET platform from Lunaphore. We also thank the Lunaphore team for their collaboration and support in successfully conducting the spatial proteomic profiling of human GBM tissues. We appreciate the support and expertise of Flagship Biosciences in the design and execution of the spatial transcriptomics profiling using the GeoMx Whole Transcriptome Atlas. Special thanks to Nicole Golden and Neil Skinner from NanoString Technologies for their assistance in troubleshooting GeoMx experiments, and to Justine Johnson from Nanolive for her support with holotomography imaging. This research was partially supported by the National Institutes of Health under grants 1R01NS121075, 1R21NS116578, and 1R21CA282979 awarded to L.P.D.

## MATERIAL AND METHODS

### EXPERIMENTAL MODEL AND STUDY PARTICIPANT DETAILS

#### Animals for *in vivo* studies

Male and female C57BL/6 J (Strain# 000664), NSG (Strain#005557), Fox Chase SCID (Strain# 001913), GREAT (Strain# 017581), CCR2-deficient (Strain #017586) and CD8KO (Strain# 002665) mice were purchased from Jackson Laboratory. The animals were housed in ventilated cages at the University of Florida’s animal facility, located in a pathogen-free environment with standard temperature and humidity conditions (22°C and 50% humidity). They were kept in a room with a 14-hour light and 10-hour dark cycle. The mice were provided with standard water and diet.

#### Intracranial implant, tumor growth monitoring and survival analysis

To assess the tumor-generating potential of TSCs and TRPCs grown in the gliomasphere assay, the tumor cells were intracranially implanted into immunocompetent C57BL/6, CD8KO (JAX colony #) mice and immunocompromised NSG and Fox-Chase SCID mice aged 7 to 15 weeks, in compliance with NIH and institutional (IACUC) guidelines for animal care and handling. The mouse colonies were housed at the University of Florida’s animal facility. Animals were anesthetized using isoflurane and administered analgesia prior to cell injection. The mice were intracranially injected with 50,000 viable cells, suspended in 2 µL of a cell suspension containing 50% methylcellulose and 50% PBS. This mixture was stereotaxically injected into the striatum of the murine brain using a sterile 5 mL Hamilton syringe fitted with a 25-gauge needle, guided by a stereotactic apparatus. Injection coordinates were set at 2.0 mm lateral to the bregma and a depth of 3.0 mm below the dura mater, as previously described ^12, 10, 97^.

The tumor volume was monitored longitudinally using an IVIS Spectrum imaging system (Xenogen, Alameda, CA, USA), measuring bioluminescence associated with luciferase activity. The mice were also observed for any neurological symptoms that might impact their quality of life. The endpoint criteria for survival analyses were based on the onset of neurological symptoms (such as decreased grip strength, ataxia, circling, paralysis, and seizure activity), changes in appearance, cranial deformity, and/or a decline in body condition score due to advanced disease progression . Predefined limits were not exceeded. The protocols (201910827, 201907966, 202100000029, and 202300000171) were reviewed and approved by the University of Florida Institutional Animal Care and Use Committee.

#### Tumor growth monitoring in CCR2KO mice and upon treatment with CSF1R inhibitor

To investigate the role of TAM recruitment within the TME as a key factor for the aggressive phenotype of TRPCs, we targeted CCR2 and CSF1R to assess their impact on tumorigenic potential and animal survival. 7-to 15 week-old mice were used for these studies. Tumor implantation, growth monitoring and survival analysis were performed as mentioned above. For the CCR2-modulation study, we compared the tumor growth kinetics and survival between TRPC and TSC tumor bearing CCR2-KO mice. As a control, we also used TRPC and TSC tumopr bearing C57BL/6 mice.

For the CSFR1-inhibition study, 7-to 15-week-old TRPC and TSC tumor bearing C57BL/6 mice were treated with *in vivo* monoclonal antibody (InVivoMAb) targeting CSFR1. The C57BL/6 mice were infused with InVivoMAb anti-mouse CSF1R antibody (BioXCell, cat#BE0213) or InVivoMAb rat IgG2a isotype control antibody, anti-trinitrophenol (BioXCell, cat#BE0089) for the control group. To overcome challenges of blood-brain barrier penetration, achieve localized depletion, and minimize systemic off-target effects, the antibody was continuously infused intratumorally for three weeks using an osmotic mini-pump. The infusion was delivered intratumorally over a 3-week period using an Alzet osmotic mini-pump (Model 1007D) at a rate of 0.5 µL per hour, with a dose of 300 µg per mouse. TRPC and TSC tumor bearing C57BL/6 mice were used as control. Tumor growth, health status, and survival were assessed as outlined previously.

#### *In vivo* fluorescence imaging to determine T-cell activation and IFN-γ production in the TME

To evaluate T-cell activation dynamics in a physiologically relevant *in vivo* context, 522×10L sorted TRPC and TSC KR158 expressing luciferase (KLuc) cells were intracranially implanted into 7-to 15-week-old GREAT mice ^98, 99, 100^, following institutional IACUC and NIH guidelines for animal care and handling. In these mice, the expression of the eYFP reporter gene is driven by the endogenous IFNγ promoter, making it a valuable tool for studying T cell activation and immune responses *in vivo*. Tumor growth was monitored via bioluminescence imaging using the IVIS system. *In vivo* fluorescence intensity was used as a surrogate marker for IFN-γ production. Fluorescence intensity was measured using a IVIS Spectrum imaging system to visualize eYFP reporter gene expression, which correlates with IFN-γ activation levels. Imaging data were normalized by comparing fluorescence intensity and tumor volume, using epi-fluorescence to measure IFN-γ production in control mice.

To further validate T-cell activation and IFN-γ production in the TME, IFN-γ levels were quantified using an ELISA. Four weeks after implantation at endpoint, brain tissues from tumor-bearing mice were harvested. The tissue was minced and incubated in papain-DNase dissociation solution (Worthington, cat# LK003150) at 37°C with constant agitation. The dissociation medium containing papain in EBSS and DNase was equilibrated with 95% O2:5% CO2 to maintain pH. After incubation, the tissue was triturated and centrifuged to collect the cell suspension. Density gradient centrifugation was then performed to separate cells from debris, and the pelleted cells were resuspended in the appropriate medium for downstream analysis. Finally, IFN-γ levels in the collected supernatant were quantified using an Invitrogen IFN-γ Platinum ELISA kit. Absorbance was measured at 450 nm using a microplate reader, and IFN-γ concentrations were determined by comparing the absorbance to a standard curve.

#### Retrospective clinical study

To perform the retrospective study (Protocol#: NH00041646; approval date: 06/12/2024 by UF Research Operations), we collected tissue samples from eligible GBM patients and performed bulk RNA sequencing (Azenta Life Sciences, Burlington, MA, USA) to assess the expression of their TRPC signature. Based on this signature expression profile, patients were stratified (into distinct groups) for further survival analysis.

##### Patient selection and data collection

The study included patients diagnosed with GBM between January 2013 and January 2024, with diagnostic codes corresponding to GBM (ICD codes: C71.0, C71.1, C71.2, C71.3, C71.4, C71.5, C70.0, C70.9, 191.0, 191.9, 191.1, 191.2, 191.3, 191.4, 191.5, 192.1), who were admitted to the Department of Neurosurgery at the University of Florida. Patient data were retrieved from deidentified electronic health records (EHRs) at Integrated Data Repository Research Services (IDR) at the University of Florida for analysis. A total of 4085 patients were identified, with 54,232 data points extracted for the analysis. Of these, survival data were available for 1408 patients, and medication data were available for 416 patients who had been prescribed lipid-lowering medications, including statins and HMG-CoA reductase inhibitors, ezetimibe, PCSK9 inhibitors, bile acid sequestrants, PPAR-alpha activators, niacin, fish oils, metformin, and various combination products. Of these, 186 patients had survival data. Survival data were defined as the number of days from the date of surgery to either the date of death or to January, 2024, whichever occurred first.

##### TRPC signature and RNA sequencing

For the subset of patients with available tissue (24 samples), RNA was extracted from freshly frozen tumor samples. Bulk RNA sequencing was performed to evaluate the expression of the TRPC signature. The raw sequencing data were processed and analyzed on the University of Florida High-Performance Computing Cluster (HiPerGator). Briefly, low-quality reads and adapter sequences in the FASTQ data were trimmed using Trim Galore (Babraham Bioinformatics). Reads with a quality score above Q30 were then aligned to the Gencode v23 human genome using RSEM ^101^. Gene expression levels were quantified, and patients were stratified based on their TRPC expression profiles.

##### Data synchronization and statistical analysis

To ensure consistency and integrity of the dataset, patient-level data were synchronized using Microsoft Excel. VLOOKUP formulas were employed to efficiently match survival data with corresponding medication records. For survival analysis, the patients were stratified based on TRPC signature and group based on median expression. The primary endpoint was OS, and Kaplan-Meier curves were constructed to visualize survival outcomes across the TRPC-stratified groups. Statistical comparisons between the survival curves were performed using the log-rank test. A significance level of p < 0.05 was considered statistically significant.

##### Ethical considerations

The study was conducted in accordance with the Declaration of Helsinki and was approved by the institutional review board (IRB) at the University of Florida. Informed consent was waived due to the retrospective nature of the study and the use of IDR as an honest broker to retrieve the deidentified patient data.

## METHOD DETAILS

### Cells

#### Glioma cell line

The murine KR158 glioma cell line (either wildtype or expressing luciferase [KLuc]) were kindly provided by Dr. Tyler Jacks at the Massachusetts Institute of Technology, Cambridge, MA, USA . The human GBM cell line used was patient-derived primary GBM Line 0 (L0) ^10^. These cells were cultured under serum-free conditions suitable for gliomasphere assays, resulting in the formation of floating spheres, as described below (in Gliomasphere Assay section).

#### Gliomasphere assay

We performed the gliomasphere assay as described in our previous study ^12^, where we established the relevance of this culture method for investigating TRPC function in a murine model of GBM. L0 and KR158 cells were cultured under serum-free conditions at 37°C with 5% CO2 in NeuroCult NS-A Proliferation Solution supplemented with 10% Proliferation Supplement (STEMCELL Technologies, Vancouver, BC, Canada; 20 ng/mL mouse EGF, and 10 ng/mL human FGF2 to stimulate murine cell proliferation. Additionally, 2ug/mL heparin was included to regulate FGF2 activity and enhance its specificity and affinity for the FGF2 receptor, thus stimulating the transduction cascade and cell proliferation. Cultures were maintained with 1% antibiotic-antimycotic. Once gliomaspheres reached a diameter of approximately 150 µm, they were enzymatically digested using Accutase for 15 minutes at 37°C. The cells were then washed, counted, and replated in fresh serum-free medium for further expansion and experimentation.

#### Isolation of TSCs and TRPCs

This gliomasphere-based paradigm was selected to enrich for distinct cellular lineages characterized by TRPC expression, as we previously reported ^11^ ^12^. These populations exhibit cancer stem cell (CSC) features, including treatment resistance, long-term persistence, and metabolic programs marked by elevated mitochondrial activity and lipid metabolism. ^11, 12^, TRPC and TSC populations grown as glioma spheres were identified and isolated from both L0 and KR158 lines based on their proliferation rates, which were assessed by their ability to retain carboxyfluorescein succinimidyl ester (CFSE), as previously described ^10^ ^102^. The CFSE/CTV fluorescence intensity decay rate over time was measured using flow cytometry. Four to seven days post-labeling, cells were grouped into CFSE/CTV^High^ (top 10%, referred to as TRPCs) and CFSE/CTV^Low^ (bottom 10%, referred to as TSCs). This gating strategy enabled the isolation of functional and phenotypic extremes with similarly sized populations, optimizing sorting time and minimizing potential fluorescence-activated cell sorting (FACS)-induced metabolic stress. By selecting these extreme fractions from the proliferation spectrum, we achieved a clear and distinct separation of TSCs and TRPCs based on their cell cycle kinetics. All experiments were carried out promptly following the FACS sorting of TRPC and TSC populations.

#### Classical adherent and serum-containing cell cultures

THP-1 and IC21 cells were cultured at 37°C with 5% CO2 in RPMI 1640, supplemented with 1% antibiotic-antimycotic and 10% or 20% FBS respectively. When the cells reached 90% confluency, they were enzymatically dissociated using Accutase for 15 minutes at 37°C, then washed and reseeded in fresh complete medium.

#### Bulk RNA-Seq sample preparation and analysis

##### Tumor dissociation for bulk RNA-seq

For all experiments involving tumor-infiltrating immune cell phenotyping and sequencing, as well as sequencing of tumor cells, tumors were isolated 3 weeks post-tumor implantation. Briefly, the right cerebral cortex (where the tumor was implanted) was excised from euthanized mice, and the tissue was processed using the Miltenyi Multi Tissue Dissociation Kit and Debris Removal Solution, following the manufacturer’s protocol. The CD45+ cells were sorted using cell FACS. Additionally, CD45 negative tumor cells were isolated from the dissociated tumor tissue for separate sequencing. The isolated cells were counted, suspended in FACS buffer, and evaluated for viability, which was typically >90% after isolation. Both the enriched CD45+ immune cells and the CD45-negative tumor cells were subsequently processed for bulk RNA sequencing.

##### Sample collection and RNA extraction

RNA extraction, library preparation, and sequencing were conducted in accordance with previously established protocols [20]. Briefly, TRPC and TSC tumors were harvested for three weeks post-implantation. RNA was simultaneously isolated from tumor cells and immune infiltrates using the Quick-DNA/RNA Miniprep Plus Kit (Zymo Research Corp).

##### Quality control and library preparation

Rigorous quality control was performed to ensure RNA purity and integrity prior to library construction. Initial RNA purity was assessed using the Nanodrop 2000 spectrophotometer (Thermo Scientific, Waltham, MA, USA), followed by agarose gel electrophoresis to examine RNA integrity and potential contamination. RNA integrity was further confirmed using the Agilent 2100 Bioanalyzer (Agilent Technologies, Santa Clara, CA, USA). mRNA was subsequently purified and randomly fragmented to facilitate cDNA synthesis. cDNA fragments of 150–200 bp were purified using AMPure XP beads (Beckman Coulter, Beverly, MA, USA) and concentrated to 1.5 ng/μL. Library quality and effective concentration were validated using the Agilent 2100 Bioanalyzer and Qubit 2.0 fluorometer (Thermo Scientific). Sequencing was carried out on the Illumina HiSeq platform following the manufacturer’s protocols.

##### Transcriptomic analysis

Multivariate principal component analysis (PCA) was performed using the FactoMineR package (version 2.11) to examine transcriptomic diversity and differentiate tumors derived from total tumor cells, TRPCs, and TSCs. Differentially expressed genes (DEGs) were identified using the R package limma (version 3.60.2), with thresholds of |logFC| > 1.5 and adjusted p-value < 0.01. Hierarchical clustering analysis was conducted using the pheatmap package (version 1.0.12).

##### Pathway enrichment and cell type deconvolution

Pathway enrichment analysis of DEGs was conducted using Gene Set Variant Analysis (GSVA) [https://doi.org/10.1186/1471-2105-14-7]. Adaptive T cell functional pathways and innate immune signatures were sourced from the Nanostring collection [https://nanostring.com/products/ncounter-assays-panels/panel-selection-tool/]. BMDM pathways were derived from [http://dx.doi.org/10.1016/j.celrep.2016.10.052], while macrophage signatures were obtained from the LM22 signature [PMC4739640]. Cell type deconvolution was performed using GSVA and ImmQuant, utilizing an immune cell gene set from Immugen [https://doi.org/10.1093/bioinformatics/btw535]. Statin sensitivity pathway analyses referenced in [https://doi.org/10.1186/s12864-021-07581-7] and [https://doi.org/10.1016/j.bbrc.2017.11.065].

##### GSEA

The assessment of gene signature enrichment was carried out using GSEA (http://www.broadinstitute.org/gsea/index.jsp. The T cell signatures, and Macrophage signature were derived from immgen.org. Nominal *p* value < 0.05 and FDR < 0.25 were applied to detect gene set enrichment across different groups.

##### Gene expression analysis using bulk RNA-seq data

FPKM values for gene expression were extracted from bulk RNA-Seq data of TSC and TRPC tumor immune infiltrates and tumors. Differential expressions of CD8, CSF1R, ABC A1, ABC G1, Iba1, CD68, CD206, Arg1, CD163, CD11b, CCR2 and FABP3 were assessed by comparing FPKM values. Statistical significance was determined using a two-tailed t-test or an appropriate non-parametric test, with a significance threshold of p < 0.05. Additionally, gene expression analysis was performed using the Subio platform, comparing RNA sequencing data for gene expression levels in immune infiltrates from TRPC tumors, TSC tumors, and normal brain tissue (non-tumor bearing). Apolipoprotein E (ApoE), the most highly expressed gene, is indicated.

##### TCR profiling

RNA sequencing was performed on CD45+ sorted cells from tumor generated by unselected Kluc cells , TRPCs, or TSCs . TCR profiling was conducted using MiXCR (version 3.0) and VDJtools (version 1.2.1), enabling alignment of sequencing reads and visualization of TCR V, D, and J gene usage. The resulting data were analyzed to identify clonotypes and their frequencies and illustrated using Circos plots.

#### Tissue dissociation and flow cytometry

Brain tissues were digested using the Multi-tissue Dissociation Kit (Miltenyl Biotec) on a gentleMACS™ Octo Dissociator with heat, followed by sample clean-up with Debris Removal Solution (Miltenyi Biotec) according to protocol as described above. Briefly, tumors were isolated 3 weeks post-tumor implantation and processed using the Miltenyi Multi Tissue Dissociation Kit and Debris Removal Solution, following the manufacturer’s protocol. The CD45+ cells were sorted using cell FACS. Samples were washed twice with cold PBS. Unstained cells were reserved for unlabeled and fluorescence-activated controls, and the cells were labeled with a live/dead dye (1:1000). Cells were washed twice in PBS and blocked for 10 minutes on ice using TruStain FcX (Biolegend) before labeling the cell surface antigens (CD8, CCR2), following the manufacturer’s recommendations. After staining, cells were filtered through a 40 µm mesh into a FACS tube (Falcon). Data acquisition was performed on the BD FACS Symphony A3, LSRII, or Canto, and analysis, including flow plots, was conducted using FlowJo version 10 (Tree Star). Doublets, dead cells, and debris were excluded, with gating based on size and granularity.

#### Exogenous lipid transfer by flow cytometry

L0 or KLuc cells were labeled with 5 µM CTV in RPMI-1640 for 15 minutes at 37°C, followed by washing with PBS as per manufacturer’s protocol. TRPCs and TSCs were isolated from these cells following the methods described above. THP-1 (for L0) and IC-21 (for KLuc) were loaded with the fluorescent lipid analog Bodipy by incubating with 10 µM Bodipy as per manufacturer’s protocol and then washed three times with PBS. The Bodipy-labeled macropahges (2.5M) were co-cultured with TRPCs and TSCs (0.5M) at a 5:1 ratio to facilitate lipid transfer and interaction for 24hrs. To study the effect of different conditions mimicking the TME, the cells were also cultured under glucose-deprived conditions with RPMI media containing 60 mg/dL of glucose and compared against normal glucose conditions of 200 mg/dL. We studied the rate of lipid efflux from the macrophages and influx into the TRPCs and TSCs.

Additionally, the rate of lipid release from the macrophages was assessed both in the presence and absence of the cancer cells to characterize the nature of lipid efflux for 72hrs. At designated time points, cells were harvested, fixed using 4% paraformaldehyde, washed, and analyzed by flow cytometry to assess lipid trafficking dynamics, using appropriate channels for Bodipy and CTV fluorescence. Data analysis was performed using FlowJo software. The rate of lipid influx (Fig. 5R) and cell viability over time (Fig. 5X,Y) were modeled using best-fit polynomial equations.

#### Serum collection and measurement of CCL2 and CCL7 by Luminex assay

Serum samples were collected from TRPC and TSC tumor-bearing mice at the experimental endpoint. Whole blood was obtained via terminal retro-orbital bleeding under deep anesthesia. Samples were allowed to clot at room temperature for 30 minutes, then centrifuged at 9,000 × *g* for 3 minutes. The resulting serum was collected, aliquoted, and stored at −80°C until analysis.

Serum levels of CCL2 (MCP-1) and CCL7 (MCP-3) were quantified using the ProcartaPlex™ Mouse Immune Monitoring Panel (48-plex; Invitrogen, Thermo Fisher Scientific), following the manufacturer’s instructions. This magnetic bead-based multiplex assay was performed using Luminex xMAP technology. Briefly, serum samples and standards were incubated with antibody-coated magnetic beads in a 96-well plate. After incubation, the plate was washed using a magnetic plate washer, followed by sequential addition of biotinylated detection antibodies and streptavidin-phycoerythrin. After a final wash, the signal was developed with reading buffer.

Fluorescence intensity was measured using a Luminex MAGPIX system, and analyte concentrations were calculated using a five-parameter logistic (5-PL) curve fit in xPONENT software. Quality control measures were included to ensure reproducibility and accuracy. The assay sensitivity was consistent with the limits of detection provided by the manufacturer.

#### Spatial proteomics analysis of immune cell populations in murine GBM TME using the NanoString GeoMx DSP analysis

Tumors were generated in C57BL/6 mice through stereotactic implantation of 50,000 viable GFP-tagged TRPCs and RFP-tagged TSCs at a concentration of 16,666 cells/ul in a 1:1 ratio, as described above. Tumor volume was monitored using IVIS, and upon reaching end point the tumors were harvested. OCT-embedded murine tumor samples were sectioned to a thickness of 8-10 µm for spatial proteomics analysis within the tumor aiming to investigate the geospatial distribution of immune infiltrates. This was performed using the GeoMx Digital Spatial Profiler (DSP) with a immune panel to analyze immune-specific markers, coupled with UV-photocleavable oligonucleotide barcodes. Barcode counting was done directly and digitally through the nCounter® Analysis System. Lineage tracing was performed using GF P and RFP expression, and the regions of interest (ROIs) were delineated based on the expression of these markers along with CD45. Thirty-five ROIs were selected from 10 sections isolated from 7 different brains. Probes from CD45-positive immune cells were collected from areas enriched in either TRPCs (GFP) or TSCs (RFP). Different tissue sections used for the study included: C6_s4.2, C7_s4.4, C7_s8.2, C8_s3.2, C8_s6.2, C9_s3.1, C11_s8.2, C12_s6.2, C13_s4.2 and C13_s4.4. C=Cassette. Here, “C” refers to the cassette, corresponding to individual mice, and “s” denotes the specific brain section (inventory ID) used for spatial profiling.

The assay used antibodies specific to immune markers, which are coupled with UV photocleavable oligonucleotide barcodes. The oligonucleotides tags are released with focused UV light from selected regions based on morphology markers. Released tags are collected and quantified with nCounter optical barcodes and counts are mapped back to the tissue location resulting in a spatially resolved profile of probes abundance. The evaluation of probe reliability and data consistency, and the identification of suitable normalization factors for accurate analysis were based on the premise that probes accurately measuring signal strength should be highly correlated with each other. More precisely, the log-ratios between them should have low SDs (similar logic to the geNorm algorithm). We applied four normalization approaches to the dataset: housekeeping gene (HK) normalization, IgG normalization (Neg), area normalization, and nuclei normalization. The agreement between IgGs and housekeeping genes suggests that normalizing with either factor minimizes artifacts in the data. Area and nuclei measurements are highly consistent with each other (SD log2 ratio of 0.31), indicating strong correlation. However, they diverge from the probe-based normalization factors, such as Neg geomean and HK geomean, implying that signal strength may not be solely dependent on area or cell count, or that these geomeans may be noisy metrics. The concordance between IgG and HK factors supports their adequacy for normalization, leading to the conclusion that area and nuclei are less reliable for signal strength measurement in this dataset. Quality control of the signals of the probes was done as follows: poorly performing probes were identified by computing the ’signal-to-background’ ratio for each target, which is the intensity of the signal divided by its IgG geomean. Probes that hovered around the background in all segments were excluded from analysis. Probes with near-background values but a few segments showing significantly higher signals were included in the analysis. Certain probes had lower background than the negative controls, and while they exhibited a wide range of values, the presence of signals well above background suggests they may provide interpretable data. This approach enabled detailed spatial analysis of 14 proteins of interest within distinct biological compartments of the tumors. The transcriptomic data for the ROIs were analyzed using Multivariate PCA with the FactoMineR package (version 2.11) to assess immune diversity expression, and the intensity of the signals associated with the markers was visualized as a heatmap.

#### Immunofluorescent staining

##### OCT sections

Brain tissues were collected from tumor-bearing mice following cardiac perfusion with cold PBS. For immunofluorescent staining, the whole brain was immersed in 4% PFA in PBS for 48 hours at 4°C, followed by washing in PBS to remove the fixative. The whole tissue was immersed in 30% sucrose in PBS overnight until the tissue sank for cryopreservation. Following this, the tissue was embedded in OCT. After embedding, the mold was quickly frozen in dry ice and quickly transferred to a -80°C freezer to solidify the OCT. For staining, 8-10 µm sections were cut using a cryostat (Microm) and mounted onto charged glass slides. When permeabilization is required, the tissue is permeabilized with 0.2% Triton X-100 in PBS for an hour, then blocked with 1% BSA-PBS at room temperature for an hour, with intermittent PBS washes (3x) after each step. The sections were then stained overnight in 0.1% Triton X-100 in PBS at 4°C with primary antibodies. The slides were washed in PBS (3x) with 0.1% Tween-20 (Sigma), then incubated with fluorescently tagged secondary antibodies for 2 hours at room temperature. Nuclei were stained with DAPI (1/1000 in PBS). Imaging was performed using a Nikon A1RMP Confocal Microscope, and the images were analyzed with Nikon NIS-Elements v4.51software and Zeiss AxioObserver D1 microscope Stitched images were acquired using a Keyence (BZ-X800 Series) imaging system, which allows for the capturing and merging of multiple image tiles to create a high-resolution, composite image.

Similarly, for OCT sections of human biopsy samples, we followed the same procedure for immunofluorescence staining.

##### FFPE sections

A portion of the human biopsy sample was fixed in 10% formalin for 24 hours at room temperature and then embedded in paraffin. The paraffin-embedded tissues were sectioned into 50 or 10 µm thick slices using a microtome and mounted onto charged glass slides depending on the experiment. Prior to immunofluorescent staining, the sections were deparaffinized in xylene and rehydrated through a graded ethanol series (100%, 95%, 70%) followed by immersion in deionized water. Antigen retrieval was carried out by heating the slides in Tris-EDTA buffer (pH 9) in a microwave for 20 minutes at 95-100C, followed by cooling to room temperature.

The following steps were performed: permeabilization, blocking, and staining with primary antibodies, secondary antibodies, and DAPI, as previously described. Imaging was performed using a Nikon A1RMP Confocal Microscope, and the images were analyzed with Nikon NIS-Elements v4.51 software. Stitched images were acquired using a Keyence (BZ-X800 Series) imaging system.

##### Oil Red O staining

OCT Brain tissue sections were prepared and stained for antibodies as above. To visualize neutral fat and fat cells, we stained using Oil Red O following manufacturer’s instructions. Briefly, the tissue was incubated in Oil Red O solution for 6 minutes followed by differentiation of the tissue using 85% Propylene Glycol in distilled water for 1 minute. After staining, sections were washed and imaged using a Zeiss AxioObserver D1 microscope for high-resolution composite images.

##### Lipid Spot staining

50um OCT brain tissue sections were stained with Lipid Spot for 10 minutes under similar conditions as above. After staining, sections were washed and imaged using a Nikon A1RMP Confocal Microscope and Keyence BZ-X800 system for high-resolution composite images.

##### 3D tissue clearing and immunolabeling

For thick brain tissue ≤2mm thickness was collected from tumor bearing mice following cardiac perfusion with cold saline. Following this, it is incubated in PBS containing 4% acrylamide (Sigma-Aldrich), 0.05% N,N’-methylenebisacrylamide (Sigma-Aldrich), 4% paraformaldehyde, and 0.25% VA-044 (TCI America). The tissue was then stored at 4 °C for 3 days to allow hydrogel permeation. After polymerization of the hydrogel at 37°C for 3 hours, the brain was sectioned into 2 mm slices and passively cleared over 3–7 days in PBS with 200 mM boric acid (Sigma-Aldrich) and 4% sodium dodecyl sulfate (Fisher Scientific), pH 8.5 at 50°C. Following the clearing process, the samples were washed in PBS with 0.1% Triton X-100 for 2 days and then immunostained at 4 °C for 2 days. The antibodies were diluted in 1% goat serum, 1% donkey serum, 0.2% Triton-X 100 and 0.05% sodium azide in PBS. As a blocking buffer, we used 5% goat serum, 5% donkey serum, 0.2% Triton-X 100 and 0.05% sodium azide in PBS. The samples were washed intermittently after each step using 0.1% Triton-X 100 in PBS. For tissue sections (≤2mm), we stained for 2 days each with primary and secondary antibodies, with 3x wash steps in between. For primary and secondary staining, tissues are placed in a 12w plate and submerged in a volume of 500uL of solution plus antibody. For washing, samples are transferred to a 50mL conical tube with 40mL of wash buffer. All staining and washing were done at RT on an orbital shaker. The samples were then whole-mounted onto slides using 62% 2,2’-Thiodiethanol (Sigma-Aldrich). Imaging was performed using a Nikon A1RMP confocal microscope, and data were analyzed with Imaris x64 v9.7.0 software.

##### 3D rendered images

The multiple sliced images obtained from the Nikon A1RMP confocal microscope (as .nd2 files) were converted into the .ims format using Imaris x64 v9.7.0 software to ensure compatibility for 3D rendering. Intensity threshold adjustments were made to precisely capture the structures of interest in each channel, optimizing the visibility of key features. The data was then segmented using the Surface tool, which was followed by refinement steps. To refine the segmentation in Imaris, thresholds were adjusted to capture only relevant intensities, and the smoothness setting was applied to reduce jagged edges. A minimum volume filter was set to exclude small noise and irrelevant structures. Manual corrections were performed using the Edit Segmentation tool to ensure accurate surface rendering. The final result was reviewed in the 3D View, with rotation, zoom, and perspective adjustments to confirm the segmentation’s precision.

To visualize overlapping structures, transparency was applied to selected objects, allowing for enhanced visualization of internal features. This method was particularly useful for examining the preferential localization of TAM s and lipid distribution within the TRPC region in comparison to the TSC region.

##### 3D lipid localization and colocalization analysis

To investigate lipid localization at the TRPC-TAM interface and explore potential lipid-related interactions within the TME, we conducted a colocalization analysis using Imaris x64 v9.7.0 software. Intensity-based colocalization was performed to assess the overlap of signal intensities between channels within defined regions of interest. The surface tool was applied to generate 3D surfaces, with intensity thresholds set to accurately capture the structures of interest in each channel. Colocalization channels were constructed by selecting specific markers for TRPC cells (FA BP7+ for humans) and TAMs (CD163+ for humans). Histogram-based thresholding was employed to refine intensity thresholds, minimizing background noise and focusing on relevant signals. Based on the generated colocalization channel, a surface was created to examine the relative distribution of lipids (Lipid Spot+) in relation to the colocalization surface, allowing us to evaluate the positioning of lipids relative to the contact zones between TRPCs and M2-TAMs within the TME.

### High-plex immunofluorescence-based analysis of the immune micro-niche in human GBM TME at a geospatial context using the COMET platform

To investigate the immune contexture within human GBM tumors, high-plex immunofluorescence imaging was performed using the COMET platform (Lunaphore Technologies, Switzerland), enabling sequential immunofluorescence assay (SeqIF, Rivest *et al*., Sci Rep. 2023; PMID: 37813886) at a single-cell resolution within the geospatial context of the tissue. FFPE biopsy samples from 5 GBM patients were used for the study: P17, P19, P21, P22, and P24. These were labeled with fluorescent antibodies targeting FABP7 to differentiate tumor regions enriched in TRPC (FA BP7 high) from TSC (i.e., non-TRPC, FA BP7 low) regions. Regions of Interest (ROIs) corresponding to TRPC and TSC areas were defined based on the FA BP7 expression pattern. A total of 39 ROIs were established from five tumor sections obtained from five different patient brains (Fig. S4a). Immune cell populations were labeled using specific markers: CD31, CD68, CD11b, CD4, HLA-DR, FoxP3, CD8 and Vimentin,. Using the COMET platform, we performed spatial mapping of immune cell populations in both tumor regions based on a customized immune panel.

HORIZON software (Lunaphore Technologies, HORIZON version 2.2.0.1) was employed to segment the nucleus, define individual cells, and delineate cellular identity based on marker expression. In brief, nuclear segmentation was conducted based on DAPI fluorescence. Cell borders were approximated by expanding nuclei border outwards with a user-defined distance, using an expansion algorithm that ensures no cell overlaps. This enabled the precise and uniform delineation of individual cells within the tissue. This was followed by feature extraction step, during which mean intensity values for each marker were calculated and were simultaneously designated spatial coordinates. This enabled the precise and uniform delineation of individual cells within the tissue. A user-defined threshold for each marker was established to capture the positive cells based on the mean signal intensity. Based on marker expression profiles, each cell was classified, assigned a biological identity. Subsequently, the HORIZON software was employed to assess the distribution of distinct cell types within the ROIs, which were categorized into TRPC and TSC regions, with the resulting data exported as CSV files for further downstream analysis. The cell type distribution was also analyzed for the percentage of cell type distribution.

### Tumor growth monitoring following treatment with FABP inhibitor

C57BL/6J mice (6-8 weeks old) were intracranially injected with 50,000 Kluc-RFP cells (16,666 cells/μL) in a 3 μL suspension. The treatment group received 250 nM of the SB-FI-26 FABP inhibitor dissolved in DMSO, while the control group received DMSO alone. Treatments were administered intracranially at a dose of 15 mg/kg on Days 0, 7, and 14 (once per week) for a total of three weeks. Tumor growth was monitored periodically using bioluminescence imaging, with signal intensity correlating to tumor volume. Tumor progression was tracked using the bioluminescent signal as a marker for tumor volume, employing the IVIS imaging system as described previously. Survival was recorded and survival curves were generated to assess the efficacy of treatment. Upon reaching the study endpoint, tissues were collected for histopathological analysis to evaluate the impact of SB-FI-26 on tumor characteristics and the immune microenvironment.

### Determination of lipid surface area in FABP inhibitor treated murine tissue

Tissues were harvested from tumor-bearing mice treated with the FABP inhibitor SB-FI-26 at study endpoint and embedded in OCT compound. Cryosections were stained using an established multiplex immunofluorescence protocol with antibodies and dyes targeting CD68, lipids (LipidSpot), along with nuclear counterstaining using DAPI, as previously described. Fluorescent imaging was performed using a Nikon A1R MP confocal microscope. To quantify lipid content, the percentage of lipid-positive area within each tissue section was calculated using FIJI (ImageJ) software. For images in which the tissue did not occupy the entire field of view, non-tissue regions were excluded by selecting the relevant area and applying the “Clear Outside” function under the “Edit” menu. The tissue area was then delineated and measured. Lipid-positive regions were identified by adjusting the threshold specific to LipidSpot staining, and the corresponding area was quantified. The lipid surface area was expressed as a percentage of the total tissue area.

### Single cell neighborhood analysis quantification (SNAQTM) in Immunohistochemical images

To perform geospatial neighborhood analysis at the single-cell level, our group developed SNAQ™, an R script designed to visualize spatial interactions in immunohistochemical images (https://github.com/AryehSilver1/SNAQ). We analyzed five biopsy specimens from GBM patients P12, P17, P19, P22 and P23. In brief, fluorescent images were segmented based on DAPI and classified using a custom CellProfiler pipeline. For the human tissues, each cell was labeled as either a TRPC (FABP7+), a M2-TAM (CD163+), or a TSC (double negative).

For mice, we analyzed endpoint tissues treated with the FABP inhibitor and compared them to controls. All images used for quantification were acquired under matched acquisition settings. Each cell in the images of mouse tumors was labeled as either a cancer cell (RFP+), a TAM (CD68+), or neither/other (RFP and CD68 double negative).

In both human and mouse tissues, each cell was further labeled as either carrying lipid droplets (LipidSpot+) or not. The classifications for each cell and their (x,y) coordinates were exported as an excel sheet and imported into SNAQ™ ^14^. Within this program, the distances between each cell and all other cells within the image were calculated. The maximum distance for cells to be considered neighbors was set to 55 microns from each cell’s center. Any cell within 55 microns of the image border was excluded from analysis, as they did not have a full neighborhood extending 55 microns in all directions. SNAQ™ also allows for the calculation of the average distance to the closest neighboring cell of a given type for any specified cell. Using this approach, we quantified the prevalence and spatial proximity of TAMs, including lipid-containing TAMs, relative to tumor cell types

### Spatial transcriptomic analysis of the whole transcriptome within the human TME using NanoString GeoMx Digital Spatial Profiling

FFPE biopsy samples from human GBM patients were sectioned to 8-10 µm thickness for Whole Transcriptome Atlas (WTA) capture, utilizing *in situ* hybridization technology with the NanoString GeoMx Digital Spatial Profiler (DSP) to characterize spatially distinct tumor regions at high resolution. Five slides containing GBM tissues from five different patients, P17, P19, P21, P22 and P24, were prepared and processed in collaboration with Flagship Biosciences (Morrisville, NC, USA) for staining and data acquisition.

Fluorescent labeling of adjacent serial sections was performed using morphological markers to distinguish TRPC (FABP7 high) and TSC (FABP7 low) areas, identify the distribution of CD45+ immune cells, and map the distribution of Nestin+ tumor cells. Using these morphological markers, 92 ROIs corresponding to TRPC and TSC regions were defined on the tissue of interest, for spatial gene expression profiling.

GeoMx RNA assays utilize in situ hybridization (ISH) probes that are attached to unique DNA indexing oligonucleotides (known as DSP barcodes) through a linker that is sensitive to UV light. Following the identification and selection of regions of interest (ROIs) on the GeoMx DSP platform, UV exposure releases the DSP barcodes, which are then collected for downstream processing. During library construction, these barcodes are labeled with information identifying their corresponding ROI and are subsequently sequenced using an Illumina sequencer. Each DNA oligo includes sequences for ROI identification (to link it back to its spatial origin), an RNA target-specific sequence (to associate it with its ISH probe), and a unique molecular identifier (UMI) used for deduplicating sequencing reads. The resulting sequence data is then analyzed and re-integrated into the GeoMx DSP software, aligning it with the tissue images and ROI selections to generate spatially-resolved profiles of RNA expressions.

Following data acquisition, quality control and normalization were conducted using the GeoMx DSP analysis suite according to NanoString’s guidelines. The quality control metrics included field of view detection percentage, binding density, nuclei count, and surface area. ROIs of different sizes were scaled by area normalization and cell counts to minimize variation between ROIs. All ROIs were selected for analysis after thorough inspection against QC parameters. Data analysis was performed using the GeoDiff and GeoMxTools packages from the Bioconductor project, with fitting done via a generalized mixed convolution model, accounting for random effects due to slides and patient ID. The normalization process deconvolved background noise (Poisson process) from a scaled gene expression (Negative Binomial process) and adjusted for variations across slides and patients within the mixed effect differential expression model.

### Deciphering Cell-Cell Communication

#### Ligand-receptor interaction using Cell Chat

To investigate the enriched ligand-receptor intercellular communication within the TME, the CellChat platform^57^ was employed. This R package infers and quantifies ligand-receptor interactions using unsupervised learning, specifically non-negative matrix factorization [http://www.cellchat.org/]. Single-cell RNA-seq data from four glioblastoma (GBM) patients ^62^ were analyzed, classifying cancer cells into TRPCs and TSCs ^13^. Immune cells were identified by CD45 expression. Using CellChat, enriched ligand-receptor interactions between TRPCs, TSCs, and immune cells were identified, revealing bidirectional signaling and key communication pathways in the TME (Wilcoxon p < 0.05).

#### Identification of enriched signaling pathways using STRING database

To build on the ligand-receptor interactions identified through CellChat, the STRINGv11 database ^63^ was employed to further investigate the enriched signaling pathways associated with the observed intercellular communication. The ligand-receptor pairs identified through CellChat were used as input for STRING, enabling the identification of significant protein-protein interactions and associated signaling pathways.

For pathway enrichment analysis, the STRING database was queried for protein-protein interactions among the identified ligands, receptors, and their downstream signaling partners. The resulting network was visualized and analyzed to determine key functional pathways enriched within the interactions between TSCs, TRPCs, and CD45+ immune cells. We selected all protein-protein interactions with an FDR (False Discovery Rate) of <0.05 and organized them in descending order of interaction strength.

### Chip Cytometry for quantitative comparison of intratissue lipid content at cellular resolution

For the comparative analysis of intratissue lipid content between TRPCs and TSCs, we employed chip cytometry, a high-throughput technique that combines tissue-specific immunohistochemical staining with flow cytometric analysis to quantitatively analyze protein expression and cellular characteristics at the single-cell level. OCT sections from six human GBM patient samples (P12, P17, P22, P23, P33, and P34) were subjected to immunohistochemistry (IHC) to detect FABP7 expression, CD163+ M2-TAMs , and lipid distribution (lipidS pot+). The IHC images were acquired and processed using QuPath v0.5.1, which performed cellular segmentation. The processed images were then converted to .fcs format for subsequent analysis using chip cytometry. For the remainder of the process, we used FlowJo v10. The gating strategy involved collecting the CD163-population and comparing lipid content between cells with the top 10% and bottom 10% mean fluorescence intensity (MFI) of FABP7, designating them as FABP7 high and FABP7 low, respectively.

### Endogenous lipid trafficking using Holotomography Imaging

To investigate intercellular endogenous lipid trafficking, we employed cell tomography holographic super-resolution 3D microscopy (3D Cell Explorer; Nanolive, Switzerland) for high-resolution imaging of lipid transfer between tumor cells and IC21 macrophages in co-culture. The Nanolive 3D Cell Explorer platform enables the measurement of intracellular density in live cells by analyzing their differential 3D refractive index. This platform uses quantitative phase imaging, which measures variations in the refractive index of cellular components without the need for staining or labeling. Lipid droplets are identified through holotomography, primarily due to their distinct optical properties. Lipid droplets have a higher refractive index than surrounding cellular components like the cytoplasm and organelles. When illuminated with polarized light, these droplets appear as bright, well-defined structures within the cell. The Nanolive platform detects this contrast in refractive index, allowing for the identification of lipid droplets based on their unique optical signature. The platform generates 3D tomograms of the cell, providing a clear view of how lipid droplets are distributed within the cell. This advanced imaging technology allows for real-time tracking and observation of the dynamic movement of lipid droplets.

4×10^4 KLuc cells were cultured in serum-containing medium as described above in LIVE96 Plates (Nanolive). 8×10^4 IC21 macrophages were co-cultured with KLuc cells for 1.5 hours in IC21 media (described above), and then imaged at time points 24h+0, 24h+15, 24h+30, and 24h+45 minutes to track lipid transfer. Real-time 3D imaging was performed to visualize endogenous lipid transfer. Sequential images were captured to observe stages of lipid transfer. The final images were processed using Imaris x64 v9.7.0 software.

### Live cell imaging of lipid trafficking using time-lapse fluorescent microscopy

To quantitatively investigate lipid trafficking between macrophages and cancer cells, we utilized a co-culture system enabling direct visualization and temporal tracking of lipid transfer. THP-1 cells, a human monocytic cell line, were cultured in RPMI-1640 medium supplemented with 10% fetal bovine serum (FBS) and differentiated into macrophage-like cells by treatment with 100 nM phorbol 12-myristate 13-acetate (PMA) for 24 hours.

Following differentiation, THP-1 cells were incubated with 10 µM of the fluorescent lipid analog BODIPY FL C16 (Invitrogen, Cat# D3922) at 37°C as per manufacturer’s protocol. After incubation, cells were washed thoroughly with PBS to remove excess unincorporated dye. In parallel, human-derived TRPCs were labeled with CTV; according to the manufacturer’s instructions, as previously described.

Labeled macrophages and CTV-labeled TRPCs were co-cultured at a 3:1 ratio in complete RPMI medium under low glucose condition of 60 mg/dL glucose. The co-cultures were maintained incubated at 37°C in a humidified atmosphere with 5% CO₂. To monitor the dynamics of lipid trafficking, cells were imaged at multiple time points using Zeiss AxioObserver D1 microscope equipped with an incubation chamber where cells were maintained under 37°C and 5% CO₂ during live-cell imaging. Imaging was performed at 0, 25, 51, and 61 minutes, as well as at extended time points of 20 hours and 44 hours post co-culture initiation. A 360-degree video showing lipid transfer from Macrophage to TRPCs was recorded using Nikon A1RMP Confocal Microscope.

A similar experiment was conducted in parallel using a murine system. Murine macrophages (IC-21 cell line) were loaded with 10 µM BODIPY FL C16 under identical conditions and co-cultured with murine-derived TRPCs that had been labeled with CellTrace Violet. The same co-culture ratio and imaging protocol were applied to assess lipid trafficking dynamics in the murine model.

### Quantifying cellular lipid droplet distance from macrophage/tumor cell boundary

Using images showing endogenous lipid trafficking from murine macrophages (IC-21 cells) murine glioma cells (KLuc) over-time obtained from the Nanolive platform (as described earlier), we quantified the distance of lipid droplets within macrophages from their cellular boundary forming the lipid-transfer interface with KLuc cells. The lipid droplets were identified by their distinctively high refractive index, exhibiting a maximum intensity of ≥120, as measured using Fiji software (PMID: 22743772). The “Straight” command in Fiji was employed to measure the distance of the lipid droplets from the macrophage membrane at four different time points: 24 hours + 0 minutes, 24 hours + 15 minutes, 24 hours + 30 minutes, and 24 hours + 45 minutes. Movement of lipid droplets were measured across the four time points, ensuring a representative sample of lipid dynamics over the observation period. This approach allowed us to track changes in the lipid droplet position relative to the macrophage boundary with cancer cells (tumor-immune cell synapse), enabling the analysis of lipid network redistribution dynamics.

### DNA-based cell assay to measure macrophage viability

To evaluate macrophage viability under glucose-depleted conditions that mimic the TME, THP-1 monocytes were first differentiated into macrophage-like cells by treatment with 100 nM phorbol 12-myristate 13-acetate (PMA) for 24 hours. Following differentiation, 1.2 × 10L cells were seeded in RPMI-1640 media containing either low glucose (60 mg/dL) or normal glucose (200 mg/dL) concentrations. Cells were incubated under these conditions for 24, 48, and 72 hours.

Cell viability was assessed using the CyQUANT™ Cell Proliferation Assay (Thermo Fisher Scientific) in accordance with the manufacturer’s instructions. After each incubation period, CyQUANT™ dye was added to the cells, and fluorescence intensity—corresponding to cellular DNA content—was measured using a Citation 3 microplate reader (BioTek). Fluorescence values were compared across conditions to determine the impact of glucose depletion on macrophage viability over time.

The details of the chemicals, antibodies, reagents, assay kits, instruments, cells, animals, software used in the study are listed in key resources table.

### Database-driven studies

#### Database-driven transcriptomic, proteomic, and survival analysis of lipid metabolism and immune signaling

##### Gene expression analysis

Human glioblastoma (GBM) biopsy samples and non-tumor control brain tissues were selected for gene expression analysis. All analyses were conducted using Gliovis Explorer, a web-based platform for visualizing and interpreting gene expression data. Transcriptomic data were retrieved from The Cancer Genome Atlas (TCGA), with a specific focus on genes involved in lipid metabolism, including *APOE*, *ABCA1*, and *ABCG1*. Gene expression profiling was performed using the Agilent 45.02 microarray platform, and expression levels in GBM samples were compared to those in non-tumor control tissues.

##### Correlation analysis between lipid metabolism and immune signaling

To investigate coordinated regulation within lipid metabolism and immune signaling pathways, two-sided Pearson’s product-moment correlation analyses were performed using bulk RNA-seq data from TCGA. Correlations were assessed between lipid trafficking genes (*ABCA1*, *ABCG1*, *APOE*) and immune-related markers (*AIF1*, *CCR2*) in GBM samples. Additional analyses examined relationships between macrophage polarization markers (*AIF1*, *CD163*, *MRC1*) and components of the CCL2-CCL7 chemokine axis (*CCL2*, *CCL7*, *CCR2*), as well as among the chemokines themselves. Parallel analyses of *ABCA1* and *ABCG1* with *AIF1* and *CCR2* were conducted to explore immune–lipid metabolic crosstalk.

##### Protein expression validation of lipid metabolism markers

To complement the transcriptomic findings and assess the spatial distribution within the TME, we examined the protein expression of *APOE*, *ABCA1*, and *ABCG1* in glioma tissues using immunohistochemistry data from the Human Protein Atlas .

##### Overall survival based on gene expression

Using GlioVis Explorer, Kaplan–Meier survival analyses were performed to assess overall survival by stratifying GBM patients into high and low expression groups based on the median expression levels of *CCL2*, *CCL7*, *AIF1*, *CD163*, *MRC1*, and *CCR2*.

##### Percentage survival based on gene signature expression

GBM patient data from the TCGA dataset were stratified into high and low expression groups based on the median value of an immune response signature (Table S8), using the PROGgeneV2 platform (https://proggene.ccbb.indianapolis.iu.edu/). Kaplan–Meier survival analysis was conducted and is presented in Fig. 7B. Differences in percentage survival between the groups were evaluated to assess the prognostic significance of immune-related gene expression in glioblastoma.

#### Statin response and signature score analysis based on retrospective data

##### Bulk-RNA sequencing data

To evaluate the relationship between TRPC score and statin sensitivity in GBM patients using the TCGA database, the patients were stratified based on their TRPC signature ^13^ and their Statin response score was determined using the Statin sensitivity signature (Table S10). We performed a two-sided Pearson’s product-moment correlation analysis to assess the strength and direction of the linear association between TRPC scores and statin response across the patient cohort.

##### Single-cell RNA sequencing data

A similar analysis was conducted using single-cell RNA-seq data. Patients were classified as TRPC or TSC based on median expression of TRPC gene signatures ^13^. A statin response score was assigned to each analyzed cell and compared between the groups.

##### Survival Analysis Based on TRPC and Immune Signatures

To evaluate the impact of statin treatment on survival, we analyzed retrospective data from a statin-treated GBM patient cohort. Patients were stratified into high and low expression groups based on the median expression of either the TRPC ^13^ (Fig.7G) or immune signature (Table S8) (Fig.7H). The differential survival outcome of these 2 subgroups (TRPC high vs TRPC low) within the statin-treated cohort was compared to the differential outcome of these 2 subgroups in the cohort of patients from the TCGA dataset (not specifically selected for statin treatment).

#### Quantification and statistical analysis

Summary of data are presented as mean+/-standard error of mean (SEM). Paired comparisons between groups were analyzed using paired t-tests for the corresponding figure legends Fig.4Q, 7I-P. For boxplots in Fig. 2D-G; 4J, K, Q; 4V, X; the boxes represent the interquartile range (from the first to the third quartile), the middle line indicates the median, and the whiskers show the minimum and maximum values within 1.5 times the interquartile range, with outliers plotted individually.

For survival analysis, we performed Kaplan–Meier survival estimation, and statistical comparisons between groups were conducted using the log-rank (Mantel–Cox) test.

For longitudinal tumor growth analysis via IVIS bioluminescence, we employed two-way ANOVA models tailored to the group structure. When group sizes were equal and data were complete across all time points, we used two-way repeated measures ANOVA. For datasets with unequal group sizes or missing values, we applied a mixed-effects model with restricted maximum likelihood (REML) to account for subject-level variability and the unbalanced design, ensuring robust analysis of tumor growth over time. A single time-point was compared between the groups using an unpaired two-tailed t-test.

All the statistics were performed using R platform and GraphPad Prism version 10.0.0.

## SUPPLEMENTAL FIGURE LEGEND

**Supplemental Figure 1.**

**A)** Graphical representation illustrating intracranial injection of TRPC and TCS KLuc cells into mice and generated tumors. The flow cytometry gating algorithm is then used to sort live immune cell population for downstream RNA sequencing analysis. (TSC n=2, TRPC n=3)

**B)** Immune pathway enrichment analysis of ImmInfs within mTRPC/mTSC tumors

**C)** Gating strategy of flow-based analysis determining infiltrating immune cells and proportion of CD8 cells within the population with negative control gating.

**D)** Circos plot from bulk TCR sequencing analysis measuring the amplification and unique arrangements of the V and J segments of the TCRβ chain showing higher frequency of specific dominant clonotype within the TSC tumor population.

**E)** Tumor burden in BL6 mice implanted with TRPCs or TSCs were quantified three weeks post injection by bioluminescence based on luciferase activity. *In vivo* fluorescent imaging of the YFP channel in both groups was acquired to serve as background signal for the imaging performed with GREAT mice.

**F-L)** FPKM values from paired-end RNA-seq analysis were used to compare **(F)** Iba1 (p<0.01), **(G)** CD68 (p<0.0001), **(H)** CD206, **(I)** Arg1, **(J)** CD163, **(K)** CD11b, **(L)** CCR2 expressions in TRPC and TSC tumors (Unpaired t-test)

**Supplemental Figure 2.**

**A-B)** Serum level expressions of CCL2 and CCL7 using ELISA. (n=3, 3; Unpaired t-test)

**Supplemental Figure 3.**

**A-B)** Correlation analysis using bulk RNA sequencing data from GBM TCGA, with Pearson’s correlation between (**A)** APOE and ABCA1, and (**B)** ABCA1 and ABCG1

**C-E)** IHC from the protein atlas of GBM patients showing tissue expression of (C) ApoE, (D) ABCA1, and (E) ABGC1

**F-I)** Correlation analysis using Pearson’s correlation coefficient between (**J)** ApoE and CCR2, **G)** ABCA1 and AIF1, **H)** ABCA1 and CCR2, and **I)** ABCG1 and CCR2 (GBM TCGA)

**J)** 3D rendering of the GBM TME showing the lipid deposition and TAM accumulation within the TRPC region. (n=5)

**Supplemental Figure 4.**

**A)** Representative image of a human GBM tissue biopsy sample, as visualized using Horizon software, with FABP7, CD68, CD8, CD3, CD11b, HLA-DR, CD4, Vimentin and CD31 channels activated to display the multi-channel image (n=5)

**B)** IHC of patient tissues stained with morphological marker FABP7 to delineate the TRPC and the TSC regions for cell phenotyping via multiseq IHC using Horizon platform (n=5)

**C)** Segmented images showing examples of the different cell types using the Horizon software: APCs, T Lymphocytes, CTLs, Th, MDSCs and TAMs. (n=5)

**D-I)** Representative images of Lipid specialized TAMs (LS-TAMs, FABP7^+^) and their close association with the TRPCs (Vimentin^+^/FABP7^+^) (n=5)

**J)** Representative IHC image of Nestin stain in Human GBM specimen (n=5)

**K)** Representative IHC image showing labeling with DAPI, FABP7 and CD163 antibodies from a serial section of J (n=5)

**L)** Cell segmentation from (S4K) using Cell profiler (n=5). “Other” signifying non-TAM, non-TRPC cells

**M)** Neighborhood analysis of the spatial distribution of the closest 10 closest M2-TAMs to TRPCs, and TSCs. Box plots comparing the distances in um. (Unpaired t-test)

**Supplemental Figure 5.**

**A)** Human biopsy tissues collected from 5 GBM tumor patients, stained with H&E for tissue morphology, while serial subsequent sections were stained for Nestin, FABP7, CD45, and DAPI for GeoMx WTA spatial transcriptomic analysis. (n=5)

**B)** Visualization of ligand-receptor pair communication network between TRPC-CD45 (STRING 12.0) (n=4)

**C)** Visualization of ligand-receptor pair communication network between TSC-CD45 (STRING 12.0) (n=4)

**D)** Bar graph showing percentage of communication contribution of ligands contributing to CSF1R activity (n=4)

**E)** Representative IHC image of human GBM biopsy sample stained for DAPI, FABP7, CD163 and LipidSpot. Inset showing segmented cells. (n=6)

**F)** Corresponding image of E converted to .fcs file arranged according to spatial coordinates (n=6)

**G, H)** Flow cytometric gating strategy for chip-cytometric analysis to assess lipid content in TRPC and TSC cells

**I)** Corresponding histograms showing lipid intensity in TRPCs and TSCs (n=6)

**J)** Corresponding IF image of (4H) (n=5)

**K)** Corresponding IF image of (4S, T) (n=5)

**Supplemental Fig. 6**

**A)** Representative holotomography images showing lipid transfer from macrophages to tumor cells through the different stages of latching, transfer, bridging and detachment (n=3)

**B)** Corresponding images of (A) depicting the peripheral cell boundaries

**C)** Representative IF image showing lipid transfer between human TRPCs (L0) and THP1 at 44hrs after co-culture (n=3)

**D)** Overlap image of C with the corresponding brightfield image

**E)** Image showing lipid transfer between murine TRPCs and IC-21 at 19hrs after co-culture

**F)** Overlap image of E with the corresponding brightfield image

**G)** Gating strategy for time-course flow cytometry to quantify differential lipid transfer between THP-1 cells and human TRPCs and TSCs from L0

**H, I)** Bar diagram of G at timepoints (H) 2hr and (I) 24hr (n=3, 3; Unpaired t-test)

**J)** Gating strategy for time-course flow cytometry to quantify differential lipid transfer between IC-21 cells, murine TRPCs, and TSCs from KR158.

**K)** Bar diagram of J at 24hr timepoint (n=3, 3; Unpaired t-test)

**L)** Lipid efflux from THP1 under different glucose conditions in absence of tumor cells. (n=6, 6; Unpaired t-test)

**M-N)** Viability of THP1 cells using Cyquant assay (n=6, 6; Unpaired t-test)

**Supplemental Fig. 7**

**A)** Representative IHC images of murine tissues treated and not treated (Control) with SB-FI-26: Nuclear stain, RFP^+^Tumor cells and CD68^+^Macrophages

**B-F)** Representative IHC images of murine GBM tissues samples showing (B) nuclear stain, (C) RFP+Tumor cells, (D) CD68, (E) Lipids, and F) Overlap image

**G)** Corresponding overlap segmented image

**Supplemental Fig. 8**

**B)** Pearson’s correlation between CCL2 and: i. AIF1, ii. CD163, iii. MRC1, iv. CCR2; CCL7, v. AIF1, vi. CD163, vii. MRC1, and viii. CCR2

**C)** Kaplan-Meier survival curves of GBM patients (TCGA) with High and Low expression of i. CCL2, ii. CCL7, iii. AIF1, iv. CD163, v. MRC1, and vi. CCR2

**D)** Correlation between OS of statin-treated UF GBM patients and TRPC score

**E)** Correlation between OS of TGCA GBM patients and TRPC score

## Notes

### Competing Interest Statement

The authors have declared no competing interest.

## REFERENCES

1 Puchalski, R. B. et al. An anatomic transcriptional atlas of human glioblastoma. Science 360, 660–663 (2018). 10.1126/science.aaf2666

2 Klemm, F. et al. Interrogation of the Microenvironmental Landscape in Brain Tumors Reveals Disease-Specific Alterations of Immune Cells. Cell 181, 1643–1660 e1617 (2020). 10.1016/j.cell.2020.05.007

3 Friebel, E. et al. Single-Cell Mapping of Human Brain Cancer Reveals Tumor-Specific Instruction of Tissue-Invading Leukocytes. Cell 181, 1626–1642 e1620 (2020). 10.1016/j.cell.2020.04.055

4 Eisenbarth, D. & Wang, Y. A. Glioblastoma heterogeneity at single cell resolution. Oncogene 42, 2155–2165 (2023). 10.1038/s41388-023-02738-y

5 Becker, A. P., Sells, B. E., Haque, S. J. & Chakravarti, A. Tumor Heterogeneity in Glioblastomas: From Light Microscopy to Molecular Pathology. Cancers (Basel) 13 (2021). 10.3390/cancers13040761

6 Mathur, R. et al. Glioblastoma evolution and heterogeneity from a 3D whole-tumor perspective. Cell 187, 446–463 e416 (2024). 10.1016/j.cell.2023.12.013

7 Xie, X. P. et al. Glioblastoma functional heterogeneity and enrichment of cancer stem cells with tumor recurrence. Neuron 112, 4017–4032 e4016 (2024). 10.1016/j.neuron.2024.10.012

8 Bonavia, R., Inda, M. M., Cavenee, W. K. & Furnari, F. B. Heterogeneity maintenance in glioblastoma: a social network. Cancer Res 71, 4055–4060 (2011). 10.1158/0008-5472.CAN-11-0153

9 Garcia-Montano, L. A. et al. Dissecting Intra-tumor Heterogeneity in the Glioblastoma Microenvironment Using Fluorescence-Guided Multiple Sampling. Mol Cancer Res 21, 755–767 (2023). 10.1158/1541-7786.MCR-23-0048

10 Deleyrolle, L. P. et al. Evidence for label-retaining tumour-initiating cells in human glioblastoma. Brain 134, 1331–1343 (2011). 10.1093/brain/awr081

11 Hoang-Minh, L. B. et al. Infiltrative and drug-resistant slow-cycling cells support metabolic heterogeneity in glioblastoma. EMBO J 37 (2018). 10.15252/embj.201798772

12 Chakraborty, A. et al. KR158 Spheres Harboring Slow-Cycling Cells Recapitulate High-Grade Glioma Features in an Immunocompetent System. Cells 13 (2024). 10.3390/cells13110938

13 Yang, C. et al. Slow-Cycling Cells in Glioblastoma: A Specific Population in the Cellular Mosaic of Cancer Stem Cells. Cancers (Basel) 14 (2022). 10.3390/cancers14051126

14 Silver, A. et al. Heterogeneity of glioblastoma stem cells in the context of the immune microenvironment and geospatial organization. Front Oncol 12, 1022716 (2022). 10.3389/fonc.2022.1022716

15 Reilly, K. M., Loisel, D. A., Bronson, R. T., McLaughlin, M. E. & Jacks, T. Nf1;Trp53 mutant mice develop glioblastoma with evidence of strain-specific effects. Nat Genet 26, 109–113 (2000). 10.1038/79075

16 Wildes, T. J. et al. Cross-talk between T Cells and Hematopoietic Stem Cells during Adoptive Cellular Therapy for Malignant Glioma. Clin Cancer Res 24, 3955–3966 (2018). 10.1158/1078-0432.CCR-17-3061

17 Liu, Y., Cai, Y., Liu, L., Wu, Y. & Xiong, X. Crucial biological functions of CCL7 in cancer. PeerJ 6, e4928 (2018). 10.7717/peerj.4928

18 Flores-Toro, J. A. et al. CCR2 inhibition reduces tumor myeloid cells and unmasks a checkpoint inhibitor effect to slow progression of resistant murine gliomas. Proc Natl Acad Sci U S A 117, 1129–1138 (2020). 10.1073/pnas.1910856117

19 Flores, C. T. et al. Lin(-)CCR2(+) hematopoietic stem and progenitor cells overcome resistance to PD-1 blockade. Nat Commun 9, 4313 (2018). 10.1038/s41467-018-06182-5

20 Stylli, S. S., Luwor, R. B., Ware, T. M., Tan, F. & Kaye, A. H. Mouse models of glioma. J Clin Neurosci 22, 619–626 (2015). 10.1016/j.jocn.2014.10.013

21 Croxford, A. L. et al. The Cytokine GM-CSF Drives the Inflammatory Signature of CCR2+ Monocytes and Licenses Autoimmunity. Immunity 43, 502–514 (2015). 10.1016/j.immuni.2015.08.010

22 Stoeckius, M. et al. Cell Hashing with barcoded antibodies enables multiplexing and doublet detection for single cell genomics. Genome Biol 19, 224 (2018). 10.1186/s13059-018-1603-1

23 Stuart, T. et al. Comprehensive Integration of Single-Cell Data. Cell 177, 1888–1902 e1821 (2019). 10.1016/j.cell.2019.05.031

24 Bungert, A. D. et al. Myeloid cell subpopulations compensate each other for Ccr2-deficiency in glioblastoma. Neuropathol Appl Neurobiol 49, e12863 (2023). 10.1111/nan.12863

25 Takacs, G. P. et al. Glioma-derived CCL2 and CCL7 mediate migration of immune suppressive CCR2(+)/CX3CR1(+) M-MDSCs into the tumor microenvironment in a redundant manner. Front Immunol 13, 993444 (2022). 10.3389/fimmu.2022.993444

26 Vakilian, A., Khorramdelazad, H., Heidari, P., Sheikh Rezaei, Z. & Hassanshahi, G. CCL2/CCR2 signaling pathway in glioblastoma multiforme. Neurochem Int 103, 1–7 (2017). 10.1016/j.neuint.2016.12.013

27 Lee, A. W. & States, D. J. Colony-stimulating factor-1 requires PI3-kinase-mediated metabolism for proliferation and survival in myeloid cells. Cell Death Differ 13, 1900–1914 (2006). 10.1038/sj.cdd.4401884

28 Al-Rifai, R., Tedgui, A. & Ait-Oufella, H. [Colony stimulating factor-1 producing endothelial cells and mesenchymal stromal cells maintain monocytes within a perivascular bone marrow niche]. Med Sci (Paris) 39, 17–19 (2023). 10.1051/medsci/2022188

29 Cersosimo, F. et al. CSF-1R in Cancer: More than a Myeloid Cell Receptor. Cancers (Basel) 16 (2024). 10.3390/cancers16020282

30 Beffinger, M. et al. CSF1R-dependent myeloid cells are required for NK[]mediated control of metastasis. JCI Insight 3 (2018). 10.1172/jci.insight.97792

31 Remmerie, A. & Scott, C. L. Macrophages and lipid metabolism. Cell Immunol 330, 27–42 (2018). 10.1016/j.cellimm.2018.01.020

32 Goossens, P. et al. Membrane Cholesterol Efflux Drives Tumor-Associated Macrophage Reprogramming and Tumor Progression. Cell Metab 29, 1376–1389 e1374 (2019). 10.1016/j.cmet.2019.02.016

33 Huang, Y. & Mahley, R. W. Apolipoprotein E: structure and function in lipid metabolism, neurobiology, and Alzheimer’s diseases. Neurobiol Dis 72 Pt A, 3-12 (2014). 10.1016/j.nbd.2014.08.025

34 Yang, L. G., March, Z. M., Stephenson, R. A. & Narayan, P. S. Apolipoprotein E in lipid metabolism and neurodegenerative disease. Trends Endocrinol Metab 34, 430–445 (2023). 10.1016/j.tem.2023.05.002

35 Phillips, M. C. Apolipoprotein E isoforms and lipoprotein metabolism. IUBMB Life 66, 616–623 (2014). 10.1002/iub.1314

36 Getz, G. S. & Reardon, C. A. Apoprotein E as a lipid transport and signaling protein in the blood, liver, and artery wall. J Lipid Res 50 Suppl, S156-161 (2009). 10.1194/jlr.R800058-JLR200

37 Hauser, P. S., Narayanaswami, V. & Ryan, R. O. Apolipoprotein E: from lipid transport to neurobiology. Prog Lipid Res 50, 62–74 (2011). 10.1016/j.plipres.2010.09.001

38 Grimm, M. O. W., Michaelson, D. M. & Hartmann, T. Omega-3 fatty acids, lipids, and apoE lipidation in Alzheimer’s disease: a rationale for multi-nutrient dementia prevention. J Lipid Res 58, 2083–2101 (2017). 10.1194/jlr.R076331

39 Wang, S., Gulshan, K., Brubaker, G., Hazen, S. L. & Smith, J. D. ABCA1 mediates unfolding of apolipoprotein AI N terminus on the cell surface before lipidation and release of nascent high-density lipoprotein. Arterioscler Thromb Vasc Biol 33, 1197–1205 (2013). 10.1161/ATVBAHA.112.301195

40 Denis, M. et al. Molecular and cellular physiology of apolipoprotein A-I lipidation by the ATP-binding cassette transporter A1 (ABCA1). J Biol Chem 279, 7384–7394 (2004). 10.1074/jbc.M306963200

41 Zhao, G. J., Yin, K., Fu, Y. C. & Tang, C. K. The interaction of ApoA-I and ABCA1 triggers signal transduction pathways to mediate efflux of cellular lipids. Mol Med 18, 149–158 (2012). 10.2119/molmed.2011.00183

42 Vaughan, A. M. & Oram, J. F. ABCG1 redistributes cell cholesterol to domains removable by high density lipoprotein but not by lipid-depleted apolipoproteins. J Biol Chem 280, 30150–30157 (2005). 10.1074/jbc.M505368200

43 Lorenzi, I., von Eckardstein, A., Radosavljevic, S. & Rohrer, L. Lipidation of apolipoprotein A-I by ATP-binding cassette transporter (ABC) A1 generates an interaction partner for ABCG1 but not for scavenger receptor BI. Biochim Biophys Acta 1781, 306–313 (2008). 10.1016/j.bbalip.2008.04.006

44 Liang, Y. et al. Gene expression profiling reveals molecularly and clinically distinct subtypes of glioblastoma multiforme. Proc Natl Acad Sci U S A 102, 5814–5819 (2005). 10.1073/pnas.0402870102

45 De Rosa, A. et al. A radial glia gene marker, fatty acid binding protein 7 (FABP7), is involved in proliferation and invasion of glioblastoma cells. PLoS One 7, e52113 (2012). 10.1371/journal.pone.0052113

46 Morihiro, Y. et al. Fatty acid binding protein 7 as a marker of glioma stem cells. Pathol Int 63, 546–553 (2013). 10.1111/pin.12109

47 Bensaad, K. et al. Fatty acid uptake and lipid storage induced by HIF-1alpha contribute to cell growth and survival after hypoxia-reoxygenation. Cell Rep 9, 349–365 (2014). 10.1016/j.celrep.2014.08.056

48 Li, B., Syed, M. H., Khan, H., Singh, K. K. & Qadura, M. The Role of Fatty Acid Binding Protein 3 in Cardiovascular Diseases. Biomedicines 10 (2022). 10.3390/biomedicines10092283

49 Hyder, A., Sheta, B., Eissa, M. & Schrezenmeir, J. Silencing the FABP3 gene in insulin-secreting cells reduces fatty acid uptake and protects against lipotoxicity. Acta Diabetol 61, 1577–1588 (2024). 10.1007/s00592-024-02325-x

50 Dulewicz, M., Kulczynska-Przybik, A., Slowik, A., Borawska, R. & Mroczko, B. Fatty Acid Binding Protein 3 (FABP3) and Apolipoprotein E4 (ApoE4) as Lipid Metabolism-Related Biomarkers of Alzheimer’s Disease. J Clin Med 10 (2021). 10.3390/jcm10143009

51 George Warren, W., Osborn, M., Yates, A. & O’Sullivan, S. E. The emerging role of fatty acid binding protein 7 (FABP7) in cancers. Drug Discov Today 29, 103980 (2024). 10.1016/j.drudis.2024.103980

52 Cordero, A. et al. FABP7 is a key metabolic regulator in HER2+ breast cancer brain metastasis. Oncogene 38, 6445–6460 (2019). 10.1038/s41388-019-0893-4

53 Kraus, N. A. et al. Quantitative assessment of adipocyte differentiation in cell culture. Adipocyte 5, 351–358 (2016). 10.1080/21623945.2016.1240137

54 Bharati, S., Anjaly, K., Thoidingjam, S. & Tiku, A. B. Oil Red O based method for exosome labelling and detection. Biochem Biophys Res Commun 611, 179–182 (2022). 10.1016/j.bbrc.2022.04.087

55 Ramirez-Zacarias, J. L., Castro-Munozledo, F. & Kuri-Harcuch, W. Quantitation of adipose conversion and triglycerides by staining intracytoplasmic lipids with Oil red O. Histochemistry 97, 493–497 (1992). 10.1007/BF00316069

56 Najem, H. et al. Central nervous system immune interactome is a function of cancer lineage, tumor microenvironment, and STAT3 expression. JCI Insight 7 (2022). 10.1172/jci.insight.157612

57 Jin, S. et al. Inference and analysis of cell-cell communication using CellChat. Nat Commun 12, 1088 (2021). 10.1038/s41467-021-21246-9

58 Yorek, M. et al. FABP4-mediated lipid accumulation and lipolysis in tumor-associated macrophages promote breast cancer metastasis. Elife 13 (2024). 10.7554/eLife.101221

59 Yang, X. et al. FABP5(+) lipid-loaded macrophages process tumour-derived unsaturated fatty acid signal to suppress T-cell antitumour immunity. J Hepatol 82, 676–689 (2025). 10.1016/j.jhep.2024.09.029

60 Hao, J. et al. Expression of Adipocyte/Macrophage Fatty Acid-Binding Protein in Tumor-Associated Macrophages Promotes Breast Cancer Progression. Cancer Res 78, 2343–2355 (2018). 10.1158/0008-5472.CAN-17-2465

61 Saha, P., Ettel, P. & Weichhart, T. Leveraging macrophage metabolism for anticancer therapy: opportunities and pitfalls. Trends Pharmacol Sci 45, 335–349 (2024). 10.1016/j.tips.2024.02.005

62 Darmanis, S. et al. Single-Cell RNA-Seq Analysis of Infiltrating Neoplastic Cells at the Migrating Front of Human Glioblastoma. Cell Rep 21, 1399–1410 (2017). 10.1016/j.celrep.2017.10.030

63 Szklarczyk, D. et al. The STRING database in 2023: protein-protein association networks and functional enrichment analyses for any sequenced genome of interest. Nucleic Acids Res 51, D638–D646 (2023). 10.1093/nar/gkac1000

64 Chitu, V., Gokhan, S., Nandi, S., Mehler, M. F. & Stanley, E. R. Emerging Roles for CSF-1 Receptor and its Ligands in the Nervous System. Trends Neurosci 39, 378–393 (2016). 10.1016/j.tins.2016.03.005

65 Easley-Neal, C., Foreman, O., Sharma, N., Zarrin, A. A. & Weimer, R. M. CSF1R Ligands IL-34 and CSF1 Are Differentially Required for Microglia Development and Maintenance in White and Gray Matter Brain Regions. Front Immunol 10, 2199 (2019). 10.3389/fimmu.2019.02199

66 Boulakirba, S. et al. IL-34 and CSF-1 display an equivalent macrophage differentiation ability but a different polarization potential. Sci Rep 8, 256 (2018). 10.1038/s41598-017-18433-4

67 Lin, W. et al. Function of CSF1 and IL34 in Macrophage Homeostasis, Inflammation, and Cancer. Front Immunol 10, 2019 (2019). 10.3389/fimmu.2019.02019

68 Flavahan, W. A. et al. Brain tumor initiating cells adapt to restricted nutrition through preferential glucose uptake. Nat Neurosci 16, 1373–1382 (2013). 10.1038/nn.3510

69 Silver, A. et al. Get to know your neighbors with a SNAQ: A framework for single cell spatial neighborhood analysis in immunohistochemical images. Comput Struct Biotechnol J 23, 4337–4349 (2024). 10.1016/j.csbj.2024.11.040

70 Lu, H. et al. Macrophages recruited via CCR2 produce insulin-like growth factor-1 to repair acute skeletal muscle injury. FASEB J 25, 358–369 (2011). 10.1096/fj.10-171579

71 Boniakowski, A. E. et al. Murine macrophage chemokine receptor CCR2 plays a crucial role in macrophage recruitment and regulated inflammation in wound healing. Eur J Immunol 48, 1445–1455 (2018). 10.1002/eji.201747400

72 El Sayed, S., et al. CCR2 promotes monocyte recruitment and intestinal inflammation in mice lacking the interleukin-10 receptor. Sci Rep 12, 452 (2022). 10.1038/s41598-021-04098-7

73 Terada, Y. et al. Tissue-resident CCR2(+) macrophage TREM-1/3 signaling is necessary for monocyte and neutrophil recruitment to injured hearts. Cell Rep 44, 115380 (2025). 10.1016/j.celrep.2025.115380

74 Slowikowski, E. et al. A central role for CCR2 in monocyte recruitment and blood-brain barrier disruption during Usutu virus encephalitis. J Neuroinflammation 22, 107 (2025). 10.1186/s12974-025-03435-1

75 Lin, E. Y., Nguyen, A. V., Russell, R. G. & Pollard, J. W. Colony-stimulating factor 1 promotes progression of mammary tumors to malignancy. J Exp Med 193, 727–740 (2001). 10.1084/jem.193.6.727

76 Pixley, F. J. & Stanley, E. R. CSF-1 regulation of the wandering macrophage: complexity in action. Trends Cell Biol 14, 628–638 (2004). 10.1016/j.tcb.2004.09.016

77 Raghu, V. K. et al. Biomarker identification for statin sensitivity of cancer cell lines. Biochem Biophys Res Commun 495, 659–665 (2018). 10.1016/j.bbrc.2017.11.065

78 Kloosterman, D. J. et al. Macrophage-mediated myelin recycling fuels brain cancer malignancy. Cell 187, 5336–5356 e5330 (2024). 10.1016/j.cell.2024.07.030

79 Sevenich, L. Lipid recycling by macrophage cells drives the growth of brain cancer. Nature 633, 777–778 (2024). 10.1038/d41586-024-02868-7

80 Jeong, S. J., Lee, M. N. & Oh, G. T. The Role of Macrophage Lipophagy in Reverse Cholesterol Transport. Endocrinol Metab (Seoul) 32, 41–46 (2017). 10.3803/EnM.2017.32.1.41

81 Wculek, S. K., Dunphy, G., Heras-Murillo, I., Mastrangelo, A. & Sancho, D. Metabolism of tissue macrophages in homeostasis and pathology. Cell Mol Immunol 19, 384–408 (2022). 10.1038/s41423-021-00791-9

82 Viola, A., Munari, F., Sanchez-Rodriguez, R., Scolaro, T. & Castegna, A. The Metabolic Signature of Macrophage Responses. Front Immunol 10, 1462 (2019). 10.3389/fimmu.2019.01462

83 Chavakis, T., Alexaki, V. I. & Ferrante, A. W., Jr. Macrophage function in adipose tissue homeostasis and metabolic inflammation. Nat Immunol 24, 757–766 (2023). 10.1038/s41590-023-01479-0

84 Minton, K. Lipid transfer from tumour-associated macrophages supports glioblastoma. Nat Rev Immunol 24, 700 (2024). 10.1038/s41577-024-01086-6

85 Xie, Y. et al. Whether statin use improves the survival of patients with glioblastoma?: A meta-analysis. Medicine (Baltimore) 99, e18997 (2020). 10.1097/MD.0000000000018997

86 Marrone, M. T. et al. Statin Use With Immune Checkpoint Inhibitors and Survival in Nonsmall Cell Lung Cancer. Clin Lung Cancer 26, 201–209 (2025). 10.1016/j.cllc.2024.12.008

87 Kristoff, T. J. et al. Statin Drugs Are Associated With Response to Immune Checkpoint Blockade in Recurrent/Metastatic Head and Neck Cancer. Cancer Med 14, e70718 (2025). 10.1002/cam4.70718

88 Kansal, V. et al. Statin drugs enhance responses to immune checkpoint blockade in head and neck cancer models. J Immunother Cancer 11 (2023). 10.1136/jitc-2022-005940

89 Liao, Y., Lin, Y., Ye, X. & Shen, J. Concomitant Statin Use and Survival in Patients With Cancer on Immune Checkpoint Inhibitors: A Meta-Analysis. *JCO Oncol Pract*, OP2400583 (2025). 10.1200/OP-24-00583

90 Cantini, L. et al. High-intensity statins are associated with improved clinical activity of PD-1 inhibitors in malignant pleural mesothelioma and advanced non-small cell lung cancer patients. Eur J Cancer 144, 41–48 (2021). 10.1016/j.ejca.2020.10.031

91 Wang, J., Liu, C., Hu, R., Wu, L. & Li, C. Statin therapy: a potential adjuvant to immunotherapies in hepatocellular carcinoma. Front Pharmacol 15, 1324140 (2024). 10.3389/fphar.2024.1324140

92 Oesterle, A., Laufs, U. & Liao, J. K. Pleiotropic Effects of Statins on the Cardiovascular System. Circ Res 120, 229–243 (2017). 10.1161/CIRCRESAHA.116.308537

93 Choudhary, A., Rawat, U., Kumar, P. & Mittal, P. Pleotropic effects of statins: the dilemma of wider utilization of statin. Egypt Heart J 75, 1 (2023). 10.1186/s43044-023-00327-8

94 Ahmadi, M. et al. Pleiotropic effects of statins: A focus on cancer. Biochim Biophys Acta Mol Basis Dis 1866, 165968 (2020). 10.1016/j.bbadis.2020.165968

95 Sodero, A. O. & Barrantes, F. J. Pleiotropic effects of statins on brain cells. Biochim Biophys Acta Biomembr 1862, 183340 (2020). 10.1016/j.bbamem.2020.183340

96 Greenwood, J., Steinman, L. & Zamvil, S. S. Statin therapy and autoimmune disease: from protein prenylation to immunomodulation. Nat Rev Immunol 6, 358–370 (2006). 10.1038/nri1839

97 Siebzehnrubl, F. A. et al. The ZEB1 pathway links glioblastoma initiation, invasion and chemoresistance. EMBO Mol Med 5, 1196–1212 (2013). 10.1002/emmm.201302827

98 Stetson, D. B. et al. Constitutive cytokine mRNAs mark natural killer (NK) and NK T cells poised for rapid effector function. J Exp Med 198, 1069–1076 (2003). 10.1084/jem.20030630

99 Reinhardt, R. L., Liang, H. E. & Locksley, R. M. Cytokine-secreting follicular T cells shape the antibody repertoire. Nat Immunol 10, 385–393 (2009). 10.1038/ni.1715

100 Reynolds, C. J. et al. Bioluminescent Reporting of In Vivo IFN-gamma Immune Responses during Infection and Autoimmunity. J Immunol 202, 2502–2510 (2019). 10.4049/jimmunol.1801453

101 Li, B. & Dewey, C. N. RSEM: accurate transcript quantification from RNA-Seq data with or without a reference genome. BMC Bioinformatics 12, 323 (2011). 10.1186/1471-2105-12-323

102 Deleyrolle, L. P., Rohaus, M. R., Fortin, J. M., Reynolds, B. A. & Azari, H. Identification and isolation of slow-dividing cells in human glioblastoma using carboxy fluorescein succinimidyl ester (CFSE). J Vis Exp (2012). 10.3791/3918

