## Supplementary material for "Treatment Resistant Persister Cells Exploit Macrophage Lipid Metabolism to Sustain Glioblastoma Growth": Suppl. figures

**Supplemental Figure 1**

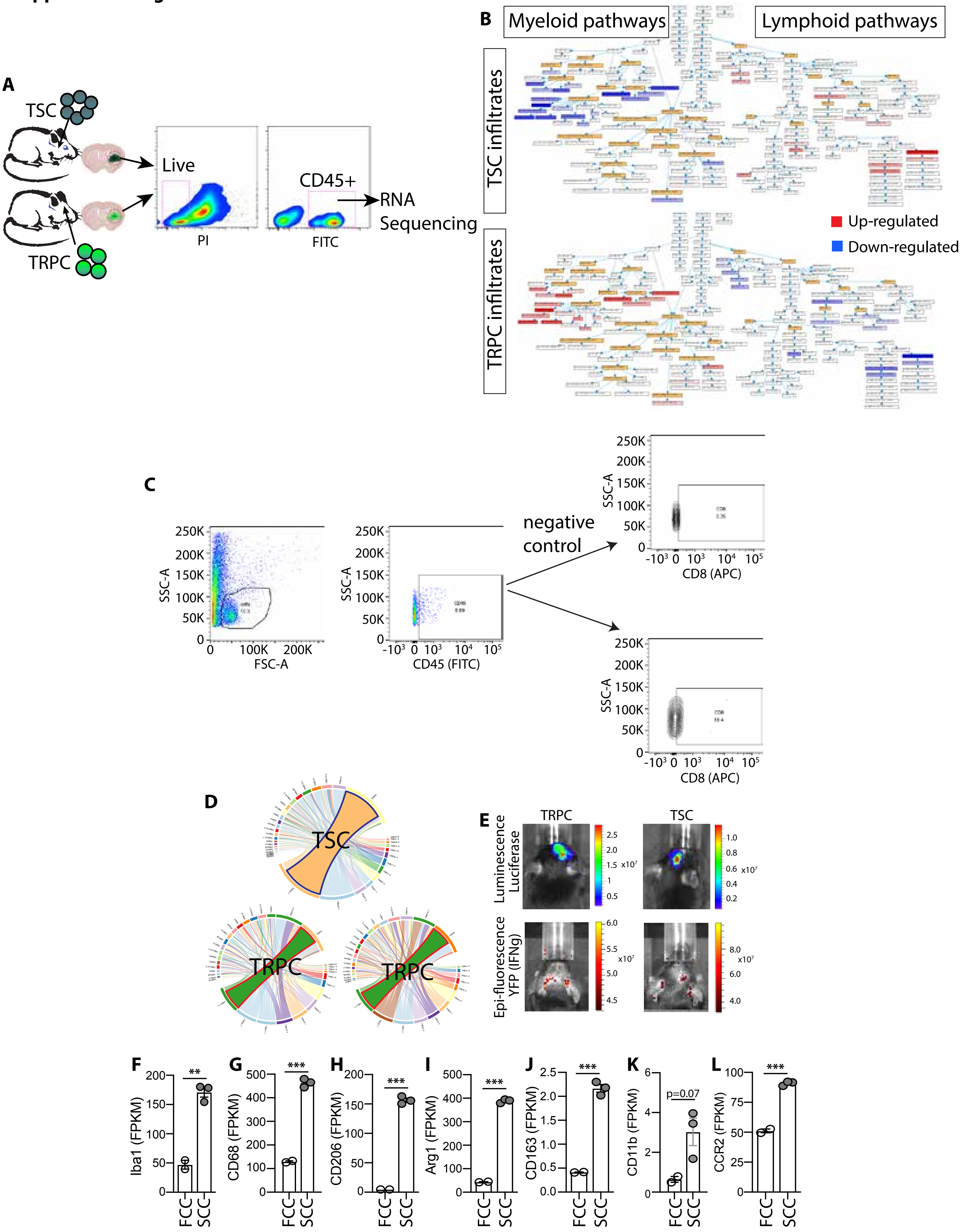

#### Supplemental Figure 2

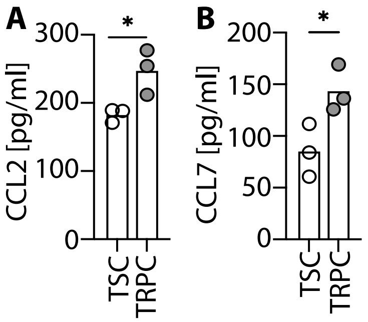

#### Supplemental Figure 3

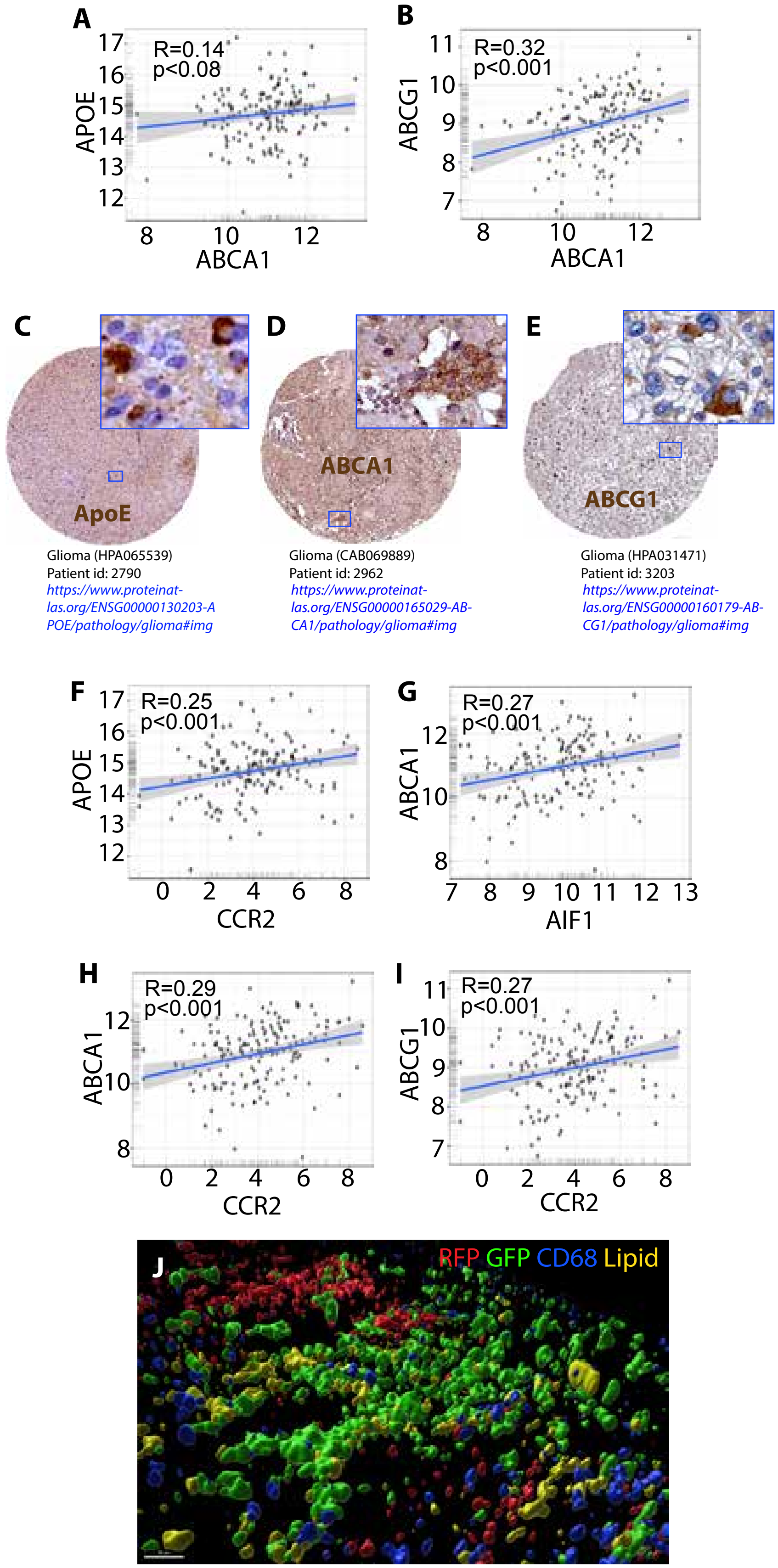

#### Supplemental Figure 4

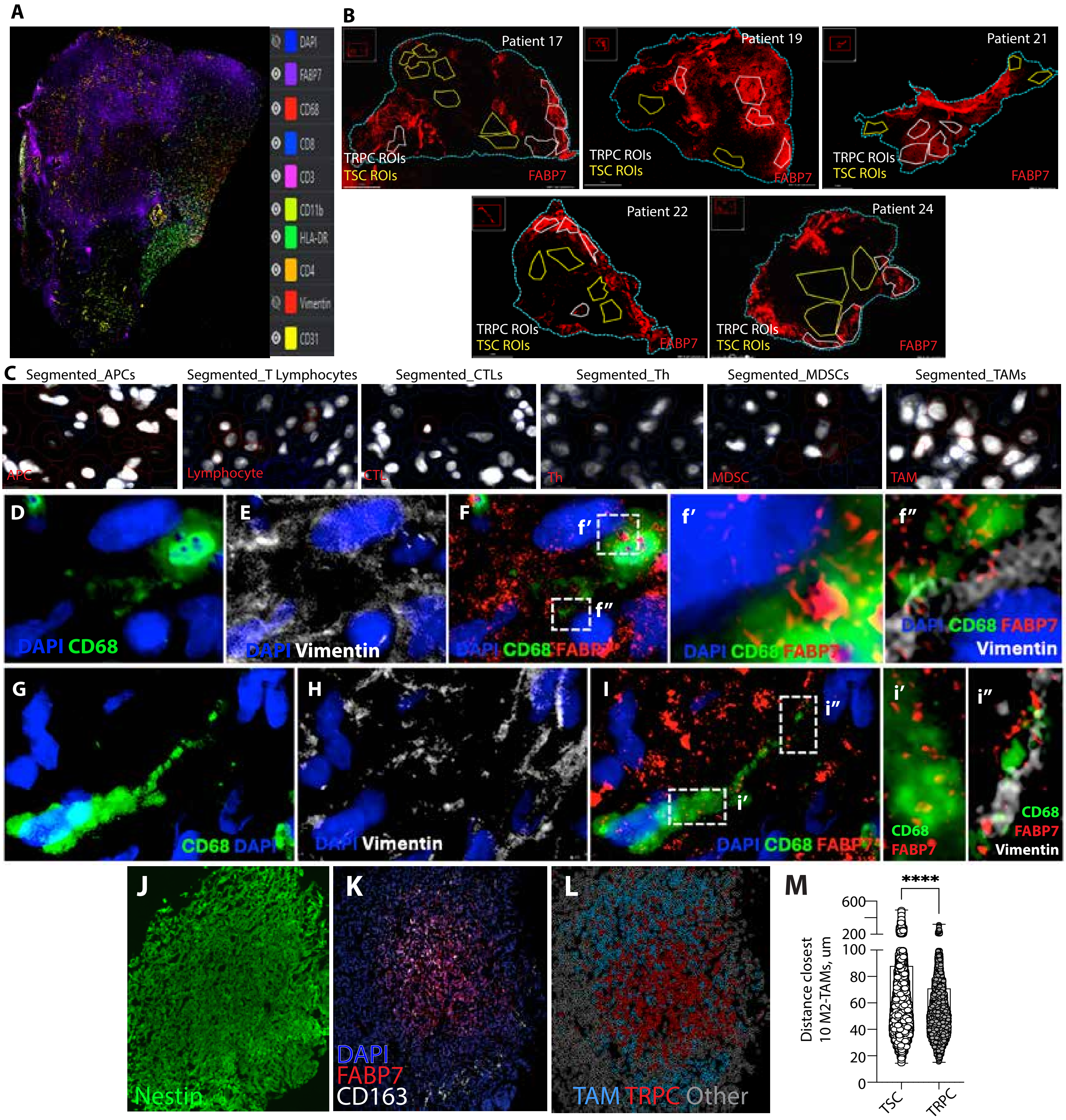

#### Supplemental Figure 5

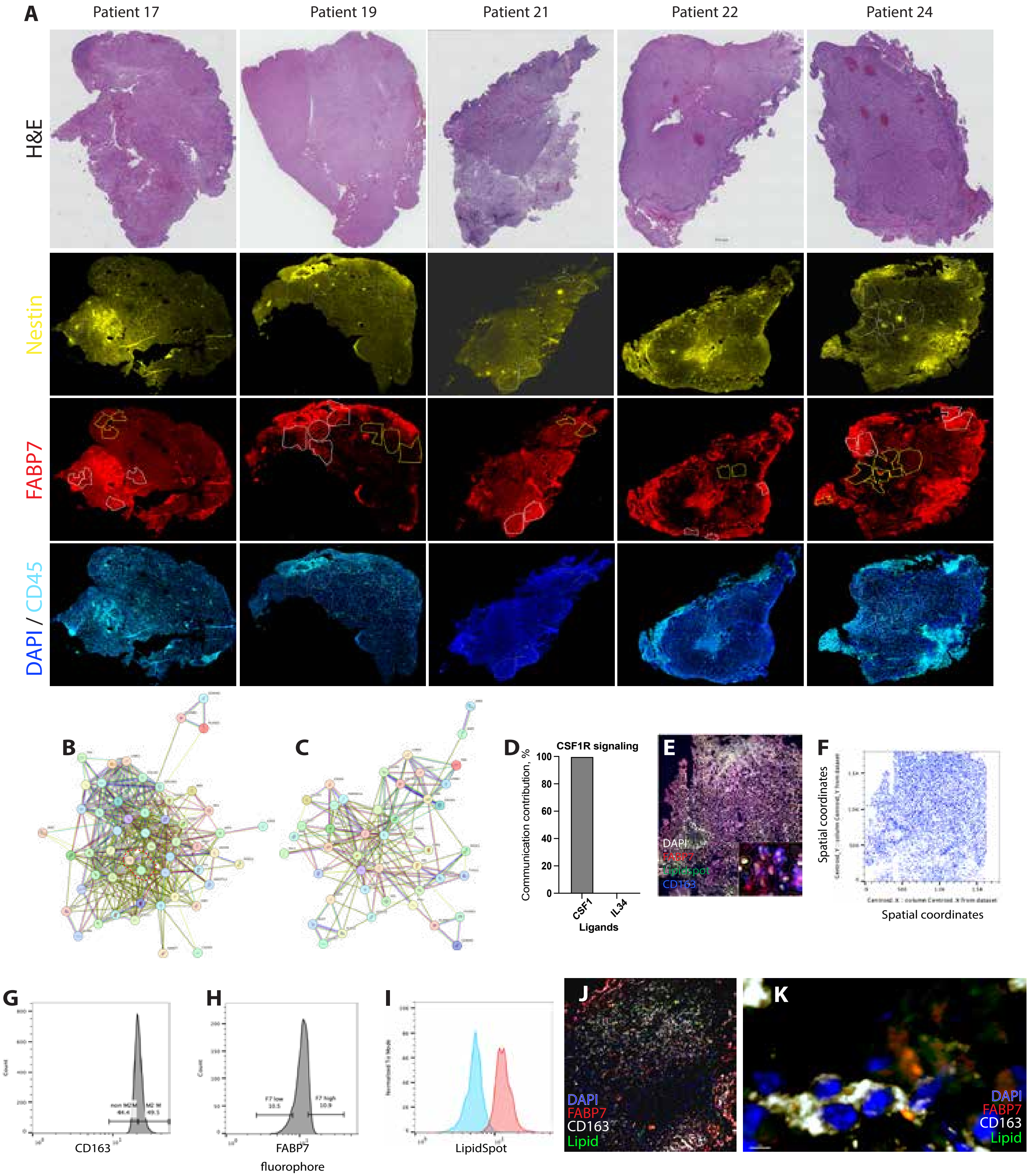

**Supplemental Figure 6**

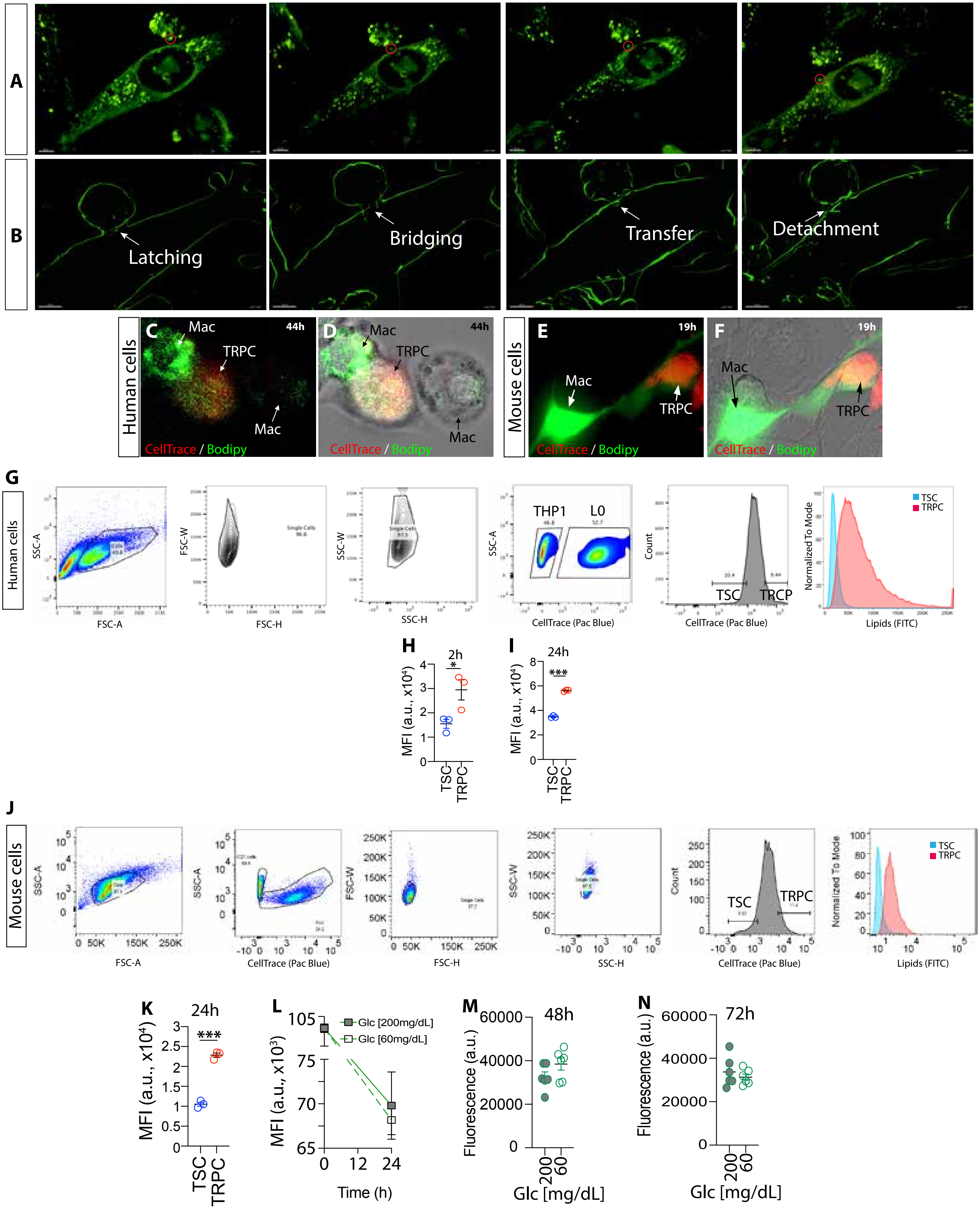

#### Supplemental Figure 7

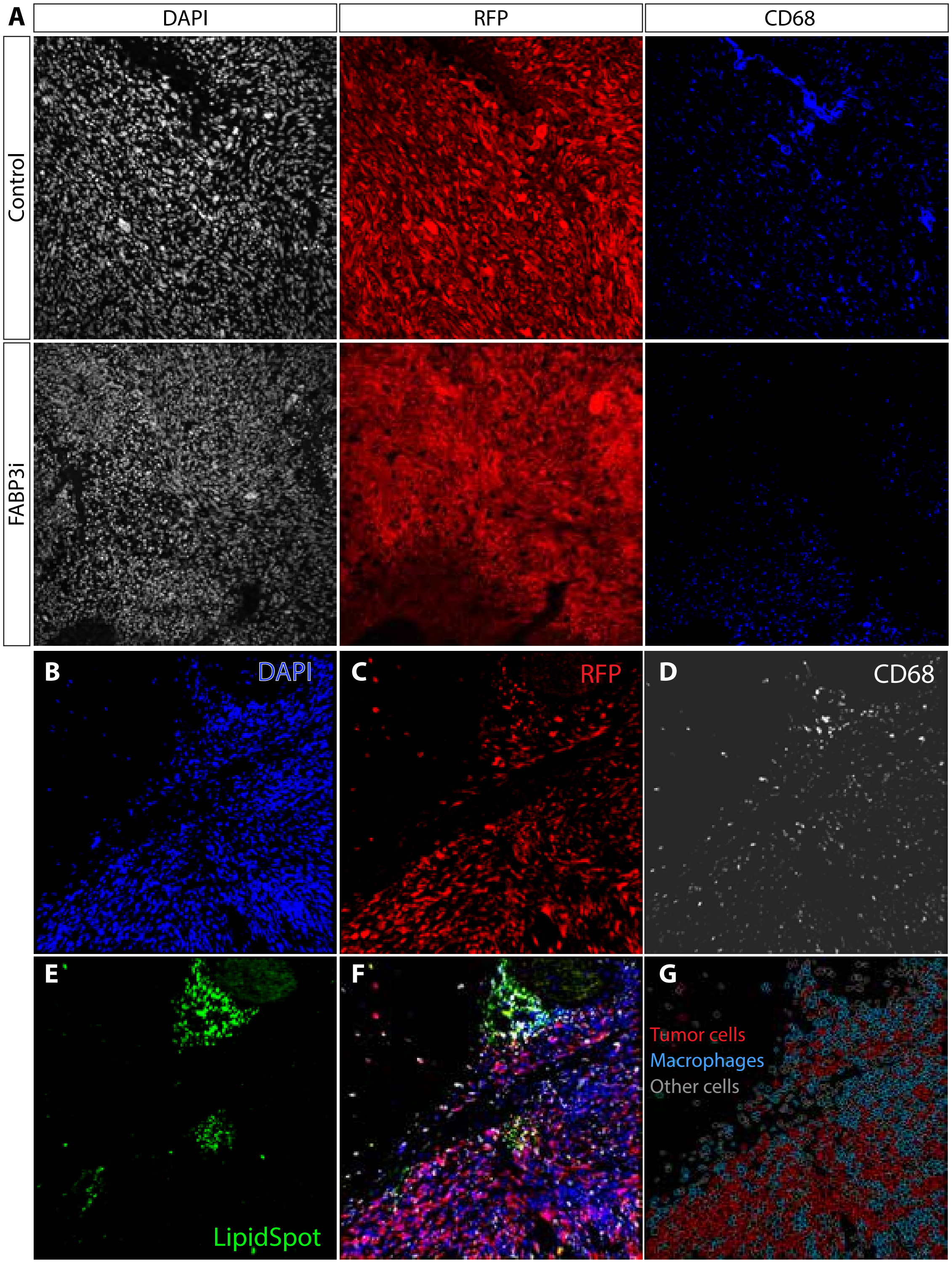

### Supplemental Figure 8

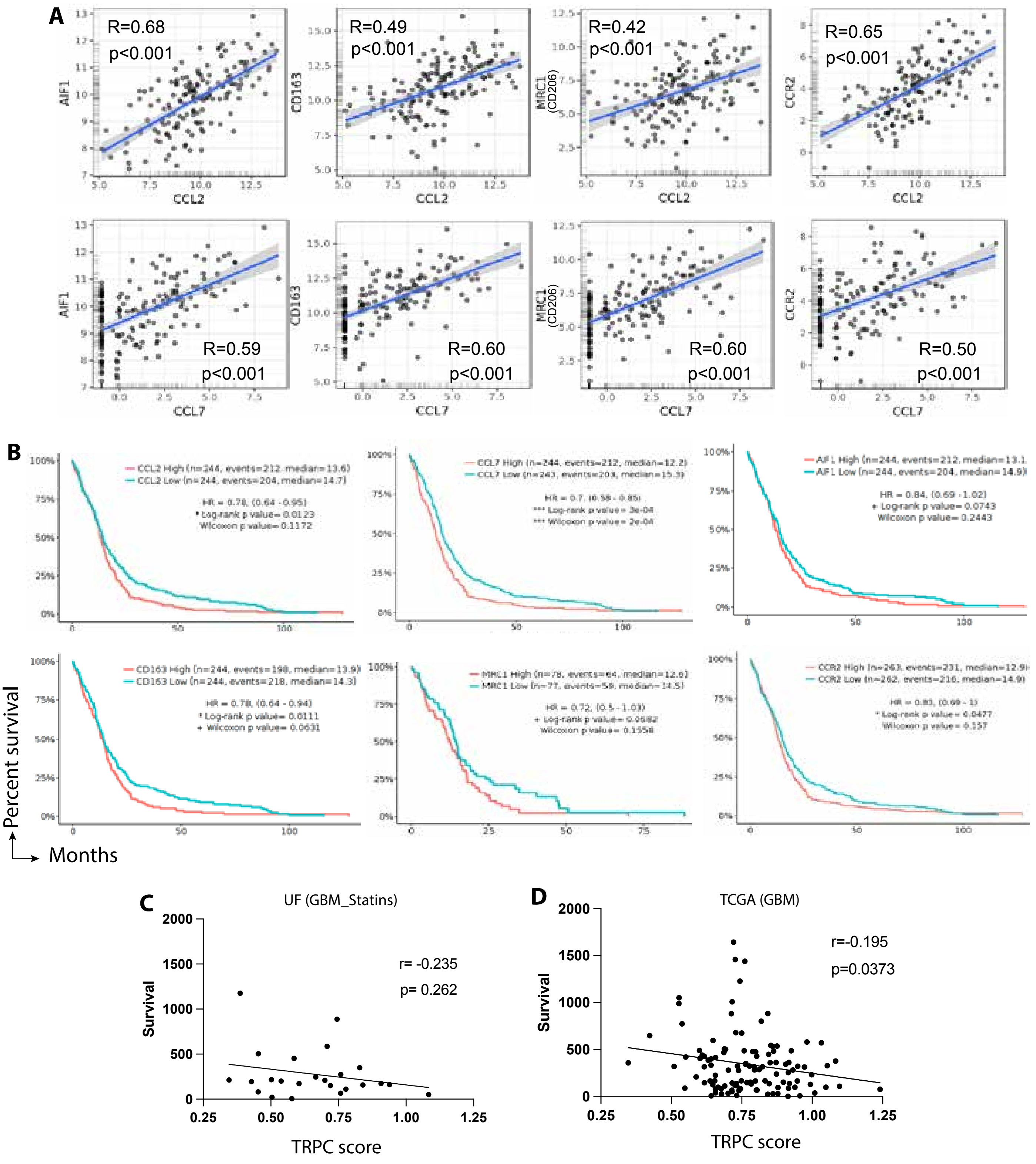
